# PI3K activation prevents Aβ42-induced synapse loss and favors insoluble amyloid deposits formation

**DOI:** 10.1101/649087

**Authors:** Mercedes Arnés, Ninovska Romero, Sergio Casas-Tintó, Ángel Acebes, Alberto Ferrús

## Abstract

Alzheimer’s disease is, to a large extent, a disease of the synapse triggered by the unbalanced amyloidogenic cleavage of the amyloid precursor protein APP. Excess of Aβ42 peptide in particular is considered a hallmark of the disease. Here we drive the expression of the human Aβ42 peptide to assay the neuroprotective effects of PI3K in adult *Drosophila melanogaster*. We show that the neuronal expression of the human peptide elicits progressive toxicity in the adult. The pathological traits include reduced axonal transport, synapse loss, defective climbing ability and olfactory perception, as well as life-span reduction. The Aβ42-dependent synapse decay does not involve transcriptional changes in the core synaptic protein encoding genes: *bruchpilot*, *liprin* and *synaptobrevin*. All toxicity features, however, are suppressed by the co-expression of PI3K. Moreover, PI3K activation induces a significant increase of 6E10 and Thioflavin-positive amyloid deposits. Mechanistically, we suggest that Aβ42-Ser26 could be a candidate residue for direct or indirect phosphorylation by PI3K. Finally, along with these *in vivo* experiments we further analyze Aβ42 toxicity and its suppression by PI3K activation in *in vitro* assays with SH-SY5Y human neuroblastoma cell cultures, where Aβ42 aggregation into large insoluble deposits is reproduced. Taken together, these results uncover a potential novel pharmacological strategy against this disease with PI3K activation as a target.

## 1. Introduction

Aβ peptide is a diagnostic biomarker of Alzheimer’s disease (AD). The peptide originates from the sequential cleavage of the Amyloid Precursor Protein (APP) by β and γ secretases in the amyloidogenic pathway, which is opposed to the non-amyloidogenic alternative (Corbett and Hooper, 2018; De Strooper et al., 2010). Through this cleavage, two predominant forms of 40 or 42 amino acid long polypeptides (Aβ40 and Aβ42) are produced (Kummer and Heneka, 2014). Only when the equilibrium between APP processing and its degradation by neprilysin, insulin-degrading enzyme or endothelin-converting enzyme (Turner et al., 2004) is imbalanced, the amyloid cascade is initiated and Aβ42 becomes synaptotoxic (Hardy and Higgins, 1992; Kummer and Heneka, 2014). The relative increase in Aβ42/Aβ40 due to excessive accumulation of β-amyloid yields protein aggregation, which involves misfolding of Aβ into soluble and insoluble assemblies. Such aggregation process is thought to be critical for AD progression, as the different Aβ assemblies differ in their toxicity (Goure et al., 2014). Whereas monomers are innocuous, when they self-associate into oligomers and pre-fibrillar aggregates they become toxic. Insoluble Aβ plaques are considered rather benign species that could serve as a sink of oligomers, although they are also a potential source of toxic aggregates (Caughey and Lansbury, 2003; DaRocha-Souto et al., 2011; Moreth et al., 2013a; Stefani, 2010). The amyloid cascade hypothesizes that plaques can sequester Aβ oligomers until reaching a physical limit after which the oligomers diffuse to the surrounding membranes and hydrophobic cell surfaces (Esparza et al., 2013).

Extracellular Aβ peptide can be detected in the cerebrospinal fluid (CSF) of AD patients (Kollhoff et al., 2018; Verberk et al., 2018). External Aβ deposits originate from APP amyloidogenic proteolysis by β-secretase, which occurs in the extracellular domain. The subsequent cleavage of the intramembranous domain by γ-secretase releases the Aβ peptide into the extracellular space or vesicle lumen (Haass et al., 2012; Shoji et al., 1992). On the other hand, intracellular Aβ accumulation has also been found inside synaptic-terminals of AD patients by high-resolution electron-microscopy (Gouras et al., 2010; LaFerla et al., 2007; Zhao et al., 2015). The nature of intracellular Aβ deposits has not yet been fully clarified, as they could be a result of endosomal membrane APP-derived Aβ, be in their way to secretion, targeted for lysosomal degradation or be a result from uptake of previously secreted Aβ oligomers. Whether the first source of toxic oligomers is the extracellular or the intracellular β-amyloid remains still an open question. Nevertheless, it is well documented that Aβ oligomers produce early synaptic alterations that eventually cause synapse loss (Li et al., 2018). At present, however, the mechanisms by which β-amyloid peptides elicit synaptotoxicity are still largely unknown.

*Drosophila* is an experimental system suitable to analyze cellular and molecular mechanisms relevant to AD. Different forms of human Aβ peptides can be genetically expressed in selected neurons in a time-controlled manner using the Gal4/UAS expression system (Brand and Perrimon, 1993). Fly models of AD follow different strategies. Some studies are based on the expression of both human APP and human BACE (β-secretase) describing the consequent Aβ plaque formation and the age-dependent neurodegeneration (Greeve et al., 2004). These events include decreased presynaptic connections, altered mitochondrial localization and reduced postsynaptic protein levels (Mhatre et al., 2014). Other models are based on the expression of human β-amyloid peptides including: Aβ40 (Iijima et al., 2004), which does not produce plaque formation but causes age-dependent learning defects; Aβ42 that elicits synaptic alterations and locomotor, survival and learning impairments (Iijima et al., 2008; Martin-Peña et al., 2017; Martín-Peña et al., 2018), and other Aβ aggregation-prone models containing human AD mutations like Aβ42-Arc (Crowther et al., 2005; Iijima et al., 2008). In addition, the 2X-UAS-Aβ42 construct, consisting in two tandem copies of the human Aβ42 fused to a secretion signal, yields stronger neurotoxic phenotypes than other single-copy models. This model shows extensive neuronal death and induces unconventional splicing of the transcription factor XBP1 (Casas-Tinto et al., 2011). Besides, 2X-UAS-Aβ42 induced neurodegeneration is rescued by XBP1 expression (Casas-Tinto et al., 2011). Thus, we have employed this construct throughout this study to generate Aβ42-dependent synaptotoxicity.

Class I phosphoinositide 3 kinase (PI3K) is involved in a signaling pathway that leads to synapse formation in *Drosophila* (Martín-Peña et al., 2006) and vertebrates (Cuesto et al., 2011). This synaptogenic pathway includes the type II BMP receptor Wit (*wishfull thinking*) and several MAPKs, but not mTOR or S6K, which are characteristic of other PI3K classical signaling pathways (Jordán-Álvarez et al., 2017). The non-canonical PI3K-mediated pathway for synaptogenesis is counterbalanced by a Mad/Medea regulated anti-synaptogenesis pathway (Jordán-Álvarez et al., 2017). In turn, inhibitors of the PI3K pathway, as GSK3-β, reduce the number of synapses (Cuesto et al., 2015). Actually, genetic manipulations of GSK3-β attenuate or suppress several AD traits in *Drosophila* (Sofola et al., 2010). Several studies have described GSK3-β activation upon Aβ expression, which can produce APP transport alterations by affecting Kinesin-1 and Dynein (Weaver et al., 2013). Consistently, GSK3-β inhibition can restore some of the Aβ toxic effects, similar to lithium or Congo-Red treatments (Crowther et al., 2004; Sofola-Adesakin et al., 2014; Sofola et al., 2010). By contrast, other reports found no restoration when GSK3-β was down-regulated in Aβ expressing neurons, claiming no Wnt-Aβ interaction in *Drosophila* AD models (Lüchtenborg and Katanaev, 2014). PI3K inhibition by downregulation of p65 regulatory subunit has also been used in Aβ, showing prevention of Aβ-induced neuronal electrophysiological defects (Chiang et al., 2010).

Synapse-promoting actions of PI3K are age-independent, as their activation at different times along adulthood generates new, supernumerary and fully functional synapses (Acebes et al., 2012, 2011; Martín-Peña et al., 2006). Recently, we reported the protective activity of PI3K over the Aβ42 toxicity caused in non-neuronal cells (Arnés et al., 2017). Unraveling the mechanisms that underlie these protective effects of PI3K would be of potential therapeutic interest, in particular if the early onset of AD could be diagnosed. Thus, considering that the phosphorylation of Aβ peptide has consequences in AD pathology (Kumar et al., 2012, 2011), we set out to investigate the potential involvement of PI3K in the mechanisms of neuroprotection against Aβ42 synaptotoxicity.

## 2 Materials and Methods

### 2.1 Fly strains

The following strains were obtained from the Bloomington Stock Center (NIH P40OD018537) (http://flystocks.bio.indiana.edu/): *elav*^*c155*^*-Gal4,* BL-458, BL-8765, BL-8760 (Lin and Goodman, 1994; Luo et al., 1994); *D42-Gal4,* BL-8816 (Chan, 2002); *Tubulin-Gal80*^*TS*^, BL-7019 (McGuire et al., 2003); *UAS-LacZ,* BL-1776 (Brand and Perrimon, 1993); *UAS GFP*^*nls*^, BL-4776 ; *UAS-PI3K*^*92D*^ (Leevers et al., 1996) and *UAS-PI3K*^*CAAX*^, BL-8294 (Parrish et al., 2009). Driver *GH298-Gal4* was a gift from Dr. Rehinard. F. Stocker (University of Fribourg) (Wong et al., 2002) whereas *krasavietz-Gal4* was provided by Dr. Gero Miessenbock (Oxford University, UK) (Dubnau et al., 2003; Shang et al., 2007). Line *UAS-Aβ42(2x)* was a gift from Dr. Pedro Fernández-Fúnez (University of Florida) (Casas-Tinto et al., 2011) and UAS-*EB1-GFP* was a gift from Dr. Melisa Rolls (Penn State University) (Rolls et al., 2007). The *UAS-Aβ42(2x)* construct contains two copies of the gene encoding the human *Aβ42* peptide which has proven effective to cause β-amyloid deposits immune-positive for 6E10 antibody (Covance). Other UAS lines include: *UAS-mTOR*, BL-7013 (Hennig and Neufeld, 2002); *UAS-bsk*, BL-9310 (Boutros et al., 1998) and *UAS-Medea*^*RNAi*^, BL-31028 (Ni et al., 2009).

### 2.2 Immunostaining

Adults were dissected and fixed with 4% formaldehyde in phosphate-buffered saline (PBS-1X) for 20 min, washed 3 times with PBS-1X 0.1% Triton-X100, and mounted in Vectashield medium with DAPI, or incubated with primary and secondary antibodies. The following antibodies and dilutions were used: mouse anti-Bruchpilot (nc82) 1:20 (DSHB); rabbit anti-HRP 1:200 (DSHB); mouse anti-Aβ42 (6E10) 1:1500 (Covance); mouse anti-phospho-Ser 1:200 (Abcam); rabbit anti-phospho-Tyr 1:200 (Abcam); mouse anti-phosphoSer8-Aβ42 1:1000 and rat anti-phosphoSer26-Aβ42 1:1000 (Dr. W. Jochen, UKB); mouse anti-β-Tubulin 1:10000 (Abcam). As secondary antibodies, we used anti-mouse Alexa 488 and anti-rabbit Alexa 568. Preparations were imaged in a Leica SP5 confocal microscope and images were processed by ImageJ. Fluorescence quantification was performed with Imaris (Bitplane) software.

### 2.3 Adult NMJ synapse analysis

Adult females of 7, 15 and 25 (± 0-3) days were dissected in order to isolate the abdominal muscles and their motor neurons. Flies were anesthetized with CO_2_ and immobilized with dissection pins in Silgar polymer plates with their dorsal side up. Flies where then submerged in a drop PBS-1X containing formaldehyde (4%), allowing the fixing process to start. A dorso-longitudinal cut was done along the abdomen and dissection pins were used to open the abdomen in a wide-open-book manner. Fat and not required tissues were removed. After a total fixation process of 20 min, the abdomens were washed with PBS-1X containing 1% Triton-X100 and transferred to 4-well plates for immunostaining.

Synapses were revealed by the primary mouse monoclonal antibody nc82, which recognizes the CAST homolog Bruchpilot protein, a constituent of the active zone (AZ) of the synapse. Anti-HRP was used to label and identify the motor neuron membrane. Samples were mounted in Vectashield medium after incubation with secondary antibodies. To quantify AZ area, we measured the total nc82 positive signal in ventral muscle packs of motor neurons in the third abdominal segment with Imaris (Bitplane) software. This estimation procedure was required as some of the experimental conditions render the nc82 signal diffuse over the motor neuron membrane as opposed to the regular appearance as single puncta (see text).

### 2.4 PI3K activating peptides

Activation of PI3K was achieved by using the peptide PTD4-PI3KAc (Cuesto et al., 2011). The peptide includes a PTD4-transduction domain that allows membrane permeabilization (Tyr-Ala-Arg-Ala-Ala-Ala- Arg-Gln-Ala-Arg-Ala), (Ho et al., 2001), fused to a phosphopeptide containing the intracellular phosphorylated SH2 domain of the PDGF receptor (Gly-Ser-Asp-Gly-Gly-pTyr-Met-Asp-Met-Ser), (Derossi et al., 1998). This peptide has been shown to induce class I PI3K activation, independently of tyrosine kinase dimerization, both *in vitro* and *in vivo* (Cuesto et al., 2011). PTD4-PI3KAc peptide was a gift from Dr. Miguel Morales (Biophysics Institute, CSIC-UPV/EHU, Spain).

### 2.5 Cell culture

SH-SY5Y human neuroblastoma cells were purchased from ATCC (ref: CRL-2266). Cells were seeded at .5×10^4^ cells/cm^2^ and used when cultures reached a 70–80% confluence. Culture media contained RPMI (Sigma-Aldrich, USA) supplemented with 0.5 mM glutamine (Sigma-Aldrich, USA), penicillin (50 mg/ml), streptomycin (50 U/ml) from Sigma-Aldrich (USA), and 10% FBS (Sigma-Aldrich, USA). Cells were serum starved and differentiated with retinoic acid (10 μM) and RPMI containing 1% FBS for 24 h prior to treatment.

### 2.6 Cell viability assay

PTD4-PI3KAc peptide was added 24 h after setting cell cultures to the wells for pre-treatment studies. Aβ42 oligomers were added to the media 24 h after the first treatment of PTD4-PI3KAc. For co-treatment experiments, PTD4-PI3KAc peptide was added at the same time as Aβ42 oligomers. Finally, for post-treatment studies, PTD4-PI3KAc peptide was added to the media 24 h after oligomer addition. PTD4-PI3KAc was added to a 50 μg/ml concentration in all cases. Concentration of Aβ42 oligomers was 18 μg/ml, 36 μg/ml or 72 μg/ml, as indicated.

### 2.7 Aβ42 intracellular aggregation assay

Aβ42 oligomers (36 μg/ml) and PTD4-PI3KAc (50 μg/ml) were added to the cell culture media. Cells were incubated with the stimuli for another 48-h period and then the coverslips followed the standard immunostaining protocol. The 6E10 immune-positive signal was analyzed in the total surface of the coverslip and quantified with Imaris (Bitplane) software.

### 2.8 Aβ42 extracellular aggregation assay

Cells were seeded and differentiated as described above. In this assay, 4 different approaches were designed: 1) Cells supernatant was transferred to a new well, with a coverslip, after 48 h of differentiation. Aβ42 oligomers were added to the supernatant and incubated for 48 h before immunostaining. This experiment was an internal control for Aβ42 aggregation. 2) Differentiated cells were treated with PTD4-PI3KAc and incubated for 48 h, the supernatant was then transferred to a new well, with a coverslip, were Aβ42 oligomers were added and incubated for another 48 h prior to immunostaining. This experiment was named PI3K pre-treatment. 3) The supernatant of differentiated cells was transferred to a new well, with a coverslip, after 48 h of differentiation. In the new well, the supernatant was treated with PTD4-PI3KAc and Aβ42 oligomers at the same time, and incubated for 48 h before immunostaining. This experiment was named PI3K co-treatment. 4) Differentiated cells were treated with Aβ42 oligomers for 48 h prior to transfer the supernatant to a new well, with a coverslip. The supernatant was then treated with PTD4-PI3KAc and incubated for another 48 h before immunostaining. This experiment was named PI3K post-treatment. The 6E10 positive dots in the total surface of the coverslip were analyzed and counted with Imaris (Bitplane) software.

### 2.9 Preparation of Aβ42 oligomers

Soluble oligomers were prepared by dissolving 0.3 mg Aβ42, previously re-solubilized in formic acid, in 200 μl of hexa-fluoro isopropanol (HFIP) for 20 min, at room temperature. 200 μl of this Aβ42 solution were added to 1 ml DD H_2_O in a siliconized Eppendorf tube. The samples were then stirred at 500 rpm using a Teflon coated micro stir bar for 8 h at 22°C for evaporation of the HFIP and progressive formation of oligomers. Before using Aβ42 oligomers for cytotoxicity assays, samples were sonicated to break and prevent incipient fibers (Kayed and Glabe, 2006).

### 2.10 Protein extraction and Western blot analysis

Ten fly heads per genotype were homogenized in 20 μl of lysis buffer containing PBS-1X, 0.05% Tween-20, 150 mM NaCl and Complete Protease Inhibitors (Roche). Equal volume of Tricine-SDS and 5% β-mercaptoethanol loading buffer was added to each sample. Samples were then boiled at 95°C for 5 min. Protein extracts were fractionated by sodium dodecyl sulfate-polyacrylamide gel electrophoresis (SDS – PAGE) in 15% Bis–Tris under reducing conditions and electroblotted into 0.22 μm nitrocellulose membranes. Membranes were then blocked in PBS containing 5% non-fat milk, and probed against Aβ42 (6E10) and β-Tubulin antibodies. To improve reactivity of 6E10 antibody, membranes were boiled for 5 min in PBS-1X before blocking. Immunoreactive bands were visualized by enhanced chemiluminescence (ECL, Amersham). Quantification of relative expression was done from three independent experiments using a loading control for normalization. The signal intensity was quantified using ImageJ (NIH) software.

### 2.11 Insoluble vs soluble protein separation

Ten fly heads were hand homogenized in 40 μl of lysis buffer and centrifuged for 1 min at 1300 rpm. The supernatant was stored as the soluble fraction. The remaining pellet was resuspended in 30 μl of PBS-1X with 1% Triton X-100 and 30 μl 70% formic acid. After sonication, vortex shaking and 800 rpm centrifugation for 1 min and the supernatant was dried with SpeedVac for 1 h. The dried pellet was finally resuspended in 40 μl of PBS-1X with 1% Triton X-100, sonicated and centrifuged again at 1300 rpm. The resulting supernatant was stored and named insoluble fraction.

### 2.12 Monomerization protocol

20 μl of monomerization buffer (9 M Urea, 1% SDS, 25 mM Tris-HCl, 1 mM EDTA) were added to total, insoluble or soluble protein fractions. The samples were then sonicated and incubated for 1 h at 55 °C, followed by centrifugation at 14000 rpm for 2 min. The supernatants were then mixed with loading buffer and boiled at 95 °C for 5 min.

### 2.13 Thioflavin-S histochemistry

7-day and 15-days (± 0-3 days) old female brains were fixed in 4% formaldehyde and permeabilized in PBS containing 0.4% Triton X-100. The brains were incubated in 50% ethanol containing 0.1% Thioflavin-S (Sigma-Aldrich) for 10 min and later washed in 50% ethanol and PBS-1X.

### 2.14 EB1-GFP live imaging

Third instar larvae were washed in PBS-1X, dried and placed in a drop of halocarbon oil 700 and 10% chloroform mixture. Dendritic arborization of dorsal cluster sensory neurons (ddA) live imaging was performed in the dorsal side of the extended larva in a spinning disk microscope. Laser intensity was maintained to 15% in all experiments. The time-lapse acquisition was done for a total time of 3 min with 2 sec intervals.

### 2.15 Negative geotaxis and lifespan assays

A total number of 30 to 48 female flies per genotype grown at 17°C were shifted to 29°C after hatching, allowing the inactivation of the Gal80^TS^ repressor and, consequently, Gal4/UAS system to be activated. Flies were separated and placed in plastic vials containing ten to fifteen of them, and kept at 29°C, 70% humidity, under a 12/12-h light/dark cycle. The counting was done at 25°C every 2-3 days in a plastic vial that was gently tapped to the bottom. The number of flies that reached the 4 cm high threshold line was counted after 10 sec of climbing; at the same time, the number of flies that reached below the 4cm criterion line and those that stayed at the bottom of the tube, were also annotated. Each counting was repeated 8 times. Lifespan assays were carried out with a similar protocol and food vials were changed every 2-3 days, and the dead flies were counted at that time. Experiments were carried out in triplicates.

### 2.16 Olfactory behavioral test and odorants

The olfactory response index, OI, was obtained in a T-maze test as described previously (Acebes et al., 2011; Acebes and Ferrús, 2001). OI score represents the number of flies counted in the odorant compartment subtracted by the number of flies trapped in the control compartment, divided by the total number of flies. OI values range from 1 (total attraction) to −1 (total repulsion) in a range of 10^−3^ to 10^−1^ concentrations, 0 value means indifference. Each data point represents the response of 350-400 individuals aged 5-7 days-old at 25°C (*GH-298 Gal4*) or 22-23 days-old at 17°C (*krasavietz-Gal4*) distributed in 14-16 replicates of 25 flies that were subjected only once to one odorant concentration choice. Experiments in which more than one-third of the flies did not make any choice were discarded. All experiments were conducted in the dark at room temperature randomizing control and experimental fly groups. No significant locomotion effects were found in any of the genotypes analyzed. The two odorants employed, Ethyl butyrate (EB) and Isoamyl acetate (IAA) were of the highest purity (Fluka, purity >95%).

### 2.17 Statistical Analysis

Data are shown as mean ± standard deviation of the mean (SD), or mean ± standard error of the mean (SEM), as indicated. Statistical significance was calculated using an unpaired Student’s two-tailed *t*-test, and an unpaired *t*-test with Welch’s correction for olfactory experiments, with significant differences between compared groups noted by \**P*≤0.05, **P≤0.01, and ***P≤0.001.

## 3 Results

### 3.1 PI3K prevents Aβ42-induced synapse loss in an age-independent manner

To determine if PI3K can rescue the loss of synapses induced by Aβ42, we overexpressed a constitutively active form of PI3K and the human Aβ42 peptide with the pan-neural *elav*-Gal4 driver. To discard an effect during development, we used the temperature sensitive Gal80^TS^ construct to activate the expression of the Gal4 system only in adult stages by a rearing temperature shift to 29°C. To estimate the number of synapses, we aged adult flies for 7, 15 and 25 days, and dissected adult abdomens to visualize the ventral longitudinal muscle (VLM) and their neuromuscular junctions (NMJs) in the third abdominal segment. Synapses were revealed by the monoclonal antibody nc82 anti-Bruch pilot (brp), whereas anti-HRP antibody was used to identify the motor neuron membrane.

At day 7 post Gal4 system activation, all genotypes had similar extents of synaptic areas, indicating that no neurodegeneration was evident when measured at the first time-point in the experimental design (**Fig. 1**). However, at day 15, Aβ42-expressing flies showed significant decrease in the total synaptic area. In addition, PI3K and Aβ42 co-expression showed also a statistically significant increase in synaptic surface compared to Aβ42-expressing flies (**Fig. 1B**). This recovery in the synaptic area indicates that the synaptogenic effect of PI3K could prevent the synapse loss induced by Aβ42 in adult motor neurons after 15 days of expression. Immune-labelled puncta in Aβ42-expressing flies showed disorganized, non-spherical positive dots of BRP signal (**Fig. 1**). This deposit-like morphology was absent in all the other genotypes. Due to this observation, the quantification of synapses was done in terms of total surface of BRP-positive signal for all the experiments, instead of the usual counting of puncta. At 25 days post-expression PI3K flies showed increased total synaptic surface, in concordance with previous published data (Martín-Peña et al., 2006). On another note, most Aβ42-expressing flies were dead and could not be analyzed, but Aβ42/PI3K flies were still alive and showed synapse areas similar to controls **(Fig.1B)**. Thus, PI3K expression prevents Aβ42 synaptotoxic effects also at later time-points.

**Figure 1.**
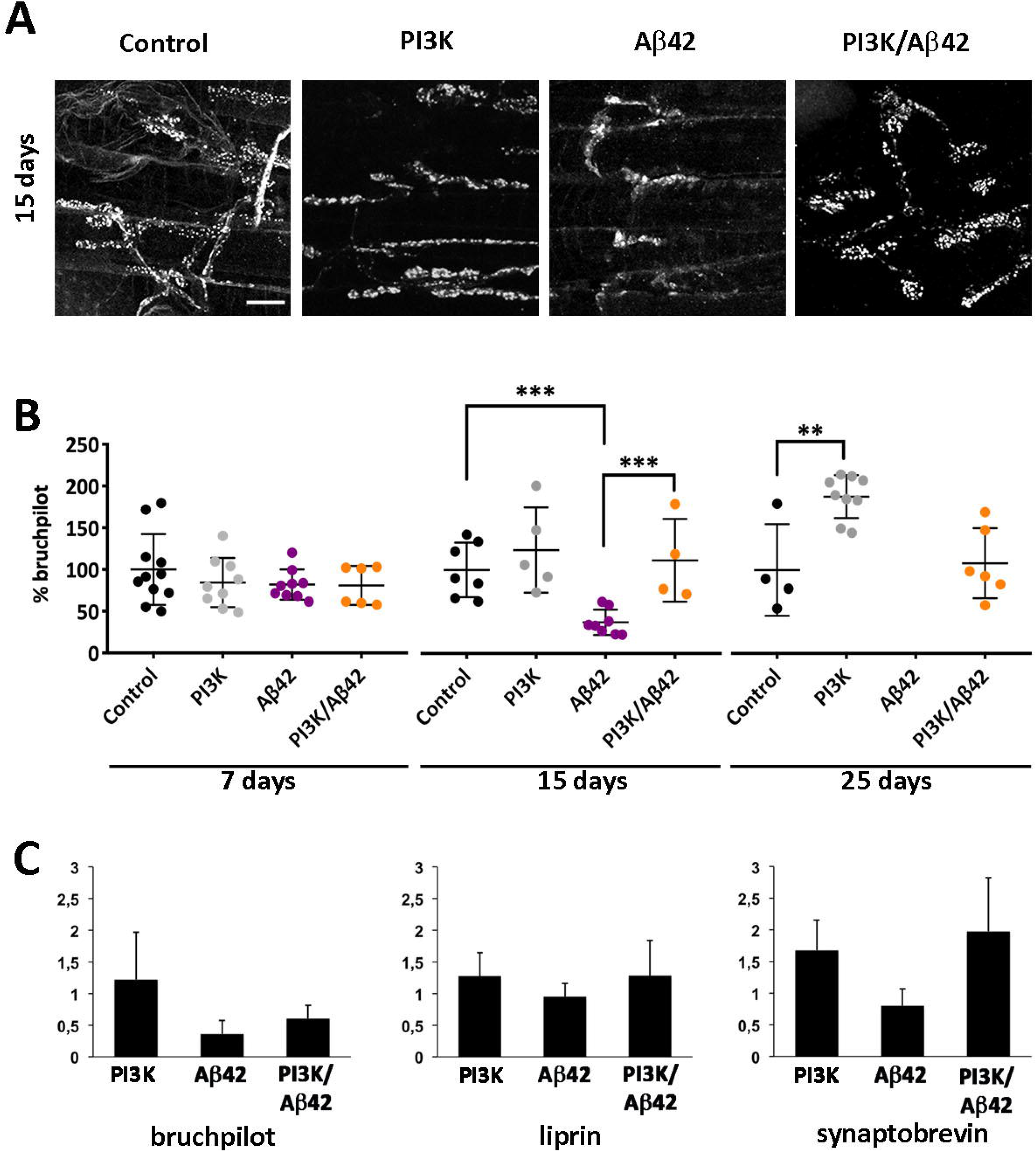
Human Aβ42 causes progressive reduction of synapses in adult motor neurons and PI3K activation suppresses this effect with no transcriptional changes in core synaptic genes. **(A)** Representative confocal images of neuromuscular junctions (NMJ) of the ventral longitudinal muscle in the third abdominal segment of adult female at 15 days post expression. Active zones monitored as nc82 immune positive area. **(B)** Time course of synaptic effects. At 7, 15 and 25 days post triggering genetic expression, the Aβ42 flies show a drastic reduction of synaptic area, while at 25 days most synapses have disappeared. Note that PI3K flies increase the synaptic signal along the three time points and the suppression effects on Aβ42 are still effective at 25 days. **(C)** Quantitative RT-PCR analysis of three genes encoding synapse proteins, *bruchpilot, liprin* and *synaptobrevin*, carried out in 15 days old adult female heads. Data are normalized to the control genotype. Histogram differences are not statistically significant. Genotypes: **Control** (*UAS-LacZ*/*elav*^*C155*^*-Gal4;* +/+; *Tub-Gal80*^*TS*^/+), **PI3K** (*UAS-PI3K*^*CAAX*^/*elav*^*C155*^*-Gal4;* +/+; *Tub-Gal80*^*TS*^/+), **Aβ42** (*elav*^*C155*^*-Gal4*/+; *UAS-Aβ42(2x)*/+; *Tub-Gal80*^*TS*^/+), **PI3K/Aβ42** (*UAS-PI3K*^*CAAX*^/*elav*^*C155*^*-Gal4; UAS-Aβ42(2x)*/+; *Tub-Gal80*^*TS*^/+). Data represent mean ± SD. Student’s t-test significances are indicated with *** p<0,001, ** p<0,01 and * p<0,05 for comparisons with the control group. Scale bar is 20μm.

To investigate a possible molecular mechanism by which PI3K rescues Aβ42-induced synapse loss, we analyzed the transcriptional status of several synaptic genes, such as *bruchpilot*, *liprin* and *synaptobrevin*, in 15 days-aged adult fly heads of the four genotypes studied here. We found no significant differences in mRNA expression in any of these genes (**Fig. 1C**), suggesting that PI3K does not restore synapses through transcriptional modulation of synaptic genes.

Taken together, we conclude that Aβ42 expression causes progressive morphological alterations and synapse loss in adult motor neurons, and that these deleterious effects can be prevented by PI3K activation in an age-independent manner without modifying gene transcription.

### 3.2 PI3K prevents the Aβ42 induced microtubule dynamics defects

Since Aβ42 can alter the neuronal cytoskeleton leading to defective neurite outgrowth (Mokhtar et al., 2013), and that this could led to deficits in intracellular transport, we tested if PI3K activation could prevent these alterations as a mechanism to protect synapses. To this end, we analyzed microtubule dynamics by live time-lapse imaging in third instar whole-mount larvae expressing the UAS-EB1-GFP construct. EB1 is a plus-end microtubule binding protein that reports microtubule dynamics, as it binds only to growing microtubules, but it detaches from shrinking microtubules resulting in dynamic GFP signals known as “comets” (Morrison et al., 1998).

We recorded EB1-GFP comets of third instar larval ddA (dendritic arborization dorsal cluster) sensory neurons (**Fig. 2A-D**). Track speed in PI3K and PI3K/Aβ42 genotypes showed similar values, being both significantly different compared to controls (**Fig. 2E,F**). In turn, track length and duration remained constant in all genotypes (**Fig. 2G,H**) indicating that the growing process of the microtubule, once started, is unaffected by Aβ42, PI3K or PI3K/Aβ42 expression. By contrast, track density was significantly decreased in Aβ42-expressing neurons, but not in PI3K and PI3K/Aβ42 expressing neurons (**Fig. 2I**), indicating that PI3K prevents the reduction in comet density caused by Aβ42 expression.

**Figure 2.**
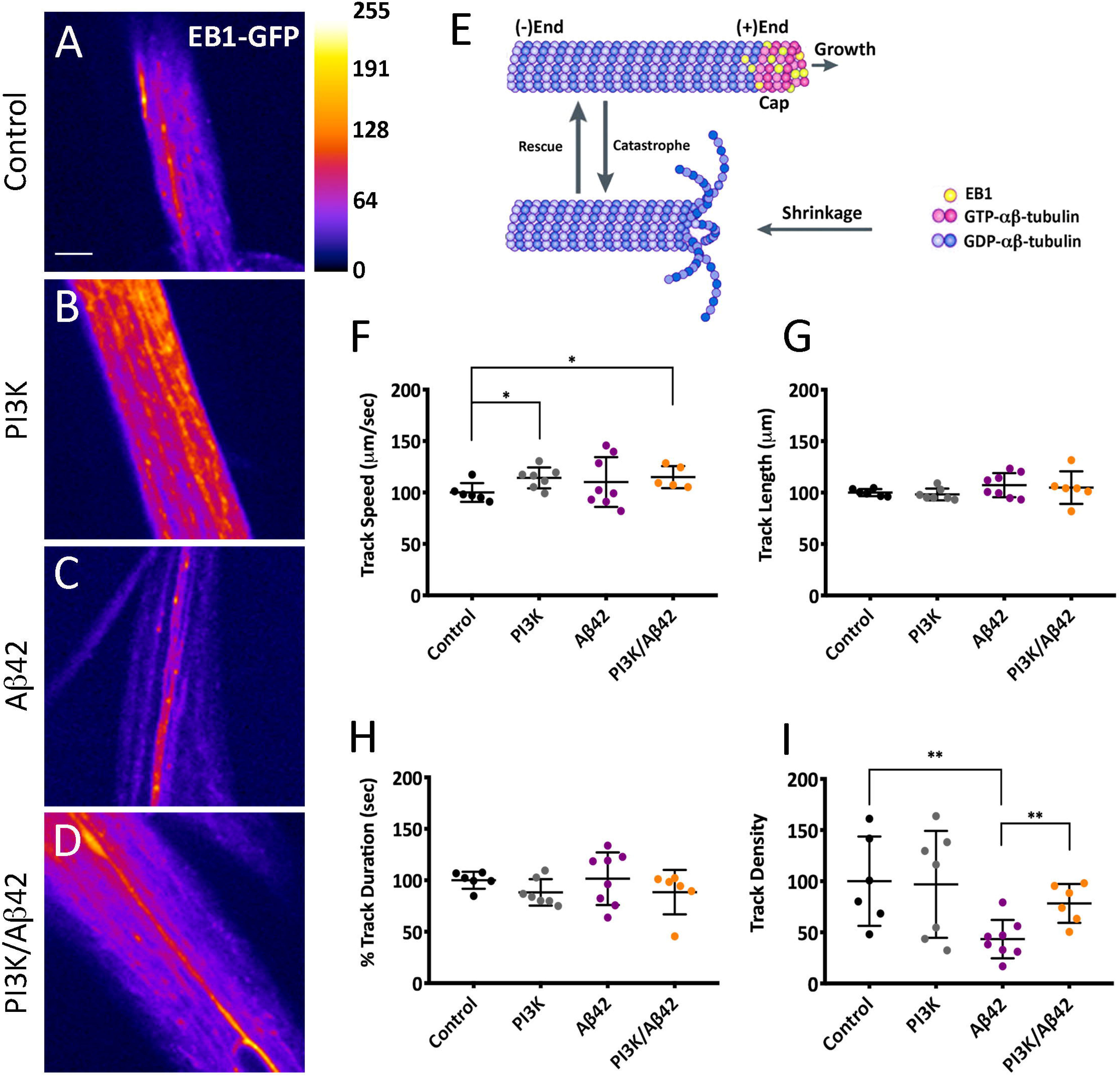
PI3K prevents Aβ42-induced microtubule dynamics deficits. **(A-D)** Still images of live recordings of EB1-GFP in ddA sensory neurons of third instar larvae. **(E)** Cartoon depicting the EB1 position in the plus-end of microtubules. **(F-I)** Quantification of GFP-tagged comet’s speed (**F**), length (**G**), duration (**H**) and density (**I**). Note the significant reduction of comet density by Aβ42 and its suppression by PI3K. Data represent mean ± SD. Student’s t-test significances are indicated with *** p<0,001, ** p<0,01 and * p<0,05 for comparisons with the control group. Scale bar is 50μm.

We concluded that Aβ42 alters microtubule dynamics by reducing the number of growing events per surface unit, which is a direct evidence of intracellular transport deficits. Interestingly, PI3K/Aβ42 neurons had the same number of growing microtubules per surface unit as control neurons, indicating that PI3K can prevent microtubule dynamics defects induced by Aβ42, thus allowing a fully functional microtubule associated intracellular transport.

### 3.3 PI3K attenuates Aβ42-induced functional deficits in locomotion, olfaction and life span

In order to test the functionality of PI3K-protected synapses under Aβ42 overexpression, we analyzed locomotion performance and olfaction in adult flies.

In locomotion assays, three different groups of flies were annotated: 1) flies that reached the 4 cm line threshold within a given time (grey histograms), 2) flies that did not reach this line but climbed along the tube (orange histograms), and 3) flies that stayed at the bottom of the tube (blue histograms) (**Fig. 3A**). Control and PI3K flies showed normal locomotion until 30 days of age, as expected for healthy flies grown at 29°C (**Fig. 3B,C**). However, Aβ42-expressing flies exhibited reduced locomotor activity as soon as 11 days of age, with a significant group of flies that stayed at the bottom of the tube during the climbing assay (black histograms) (**Fig. 3D**). The total percentage of flies that did not climb at all increased progressively as the flies aged, and this proportion of flies was higher than 50% at 18 days of age. When combined with Aβ42, PI3K overexpression delayed the appearance of the non-climbing phenotype to 15 days, which reached 50% of the total population at 25 days of age, 7 days later than the Aβ42 expressing flies (**Fig. 3E**).

**Figure 3.**
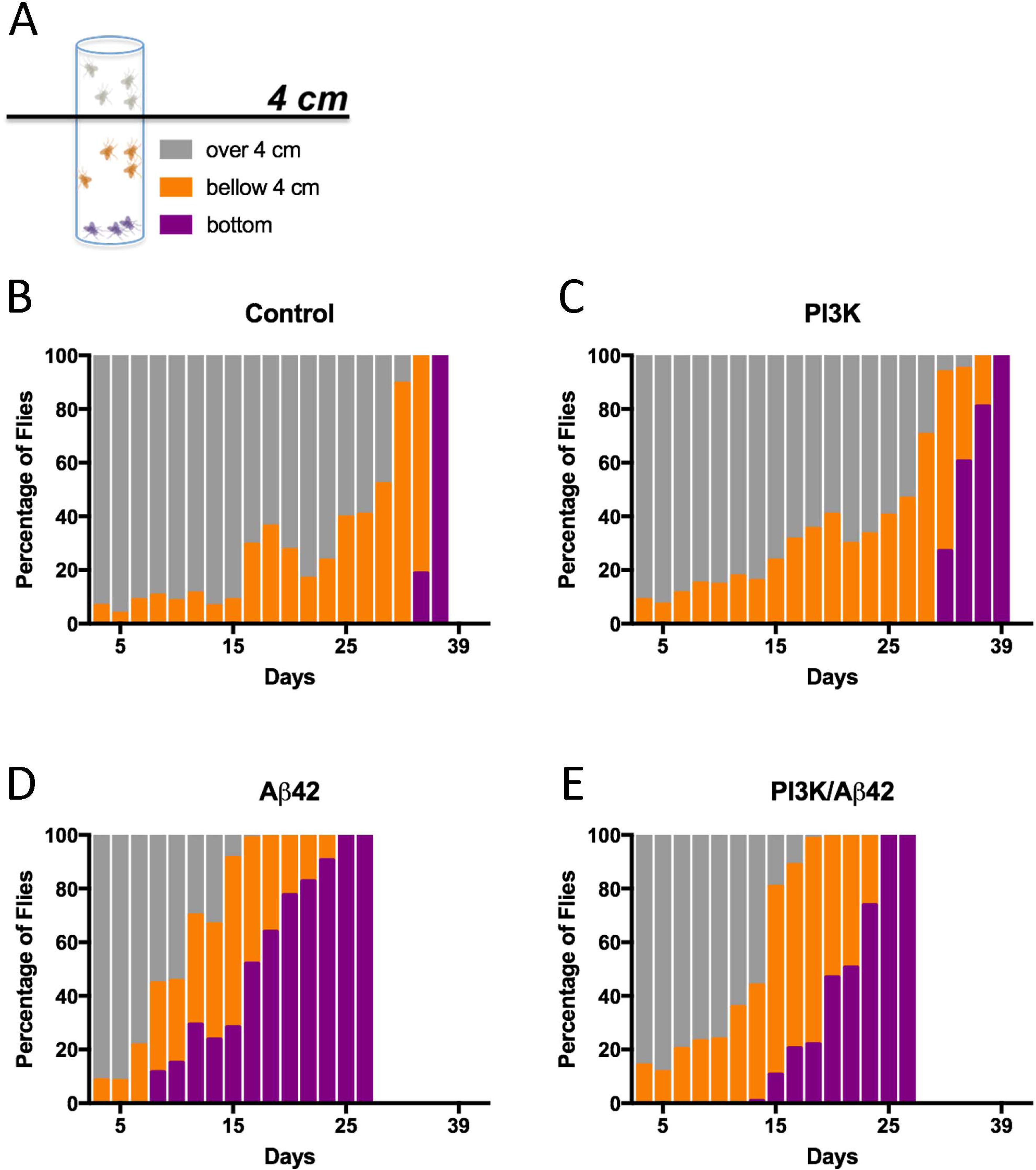
Functional effects of synapse loss and their rescue by PI3K. **(A)** Cartoon representing the three color-coded fly populations considered in the negative geotaxis assay. **(B-E)** Histograms show normalized values of flies climbing to the top of the tube (grey), not reaching the 4 cm line (orange), and not climbing (purple). Numerical data are shown in **Supp. Table S1**.

To analyze potential changes in olfactory perception, we evaluated the odorant choice index in a 10^−3^ to 10^−1^ (v/v) concentration range for two different volatiles: ethyl butyrate (EB) and isoamyl acetate (IAA) in a *GH298-Gal4* fly line (**Fig. 4** and **Suppl. Fig. S1**). This line expresses the Gal4 driver mainly in 30-32 inhibitory local interneurons of each antennal lobe, the first olfactory neuropil in the brain insect (Acebes et al., 2011; Ng et al., 2002). In the olfaction experiments we used a UAS-PI3K construct, instead of the constitutively active form UAS-PI3KCAAX, in order to test if the protective effects by PI3K would be different depending on the inducible versus the constitutively active forms of this kinase. First, we addressed the effects of Aβ42 introgression in the *GH298-Gal4* domain by using two genotypes: *UAS-Aβ42* and *UAS-Aβ42/UAS-GFP*^*nls*^. Flies overexpressing Aβ42 in either combination showed more negative olfactory indexes (**Fig. 4A**). However, the simultaneous co-expression of PI3K restored normal olfactory responses. In turn, flies expressing PI3K exhibited comparable olfactory responses with respect to control. Equivalent results were obtained in olfaction assays using isoamyl acetate, IAA, as odorant stimulus (**Suppl. Fig. S1**).

**Figure 4.**
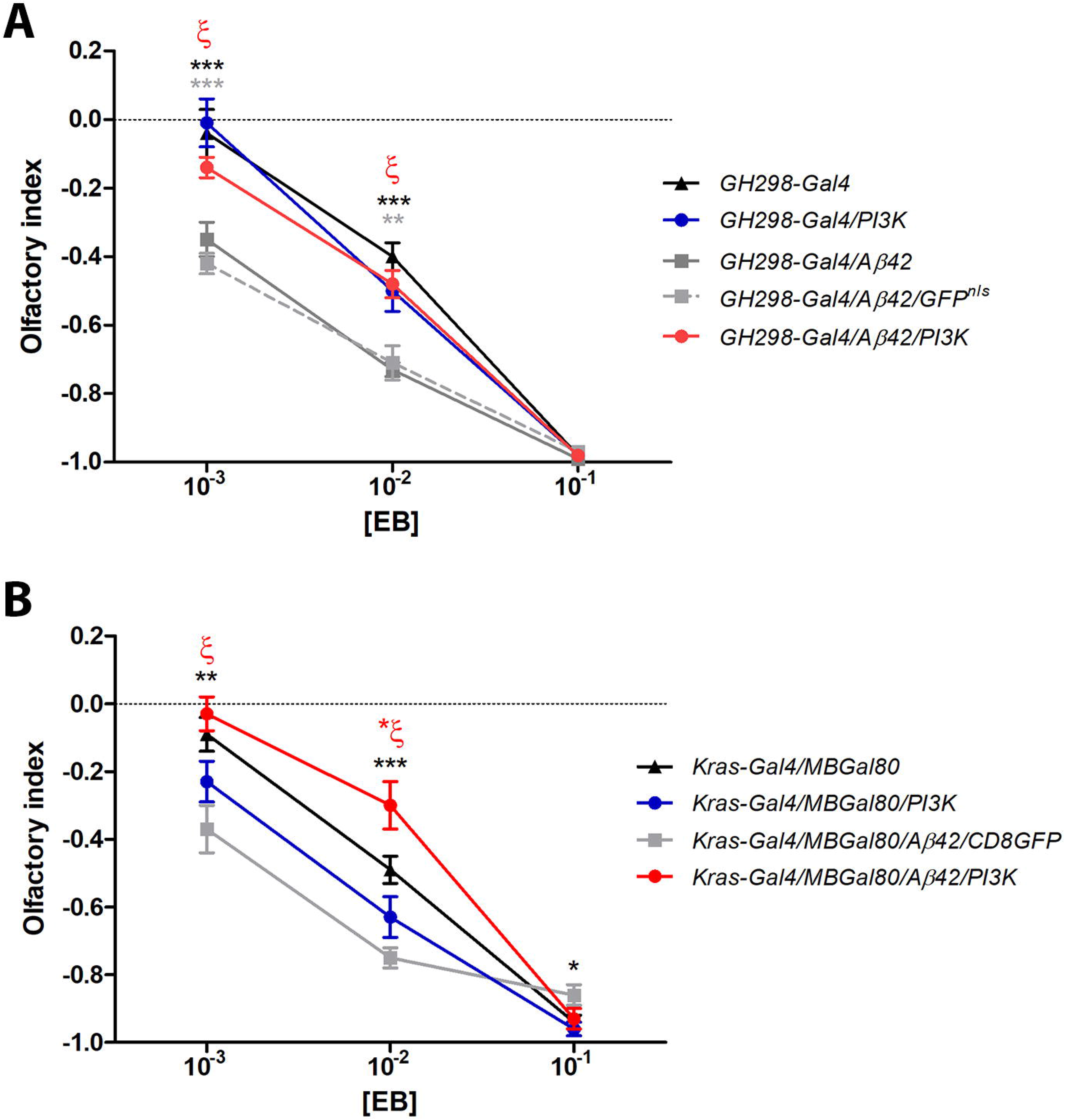
Olfactory perception assays. **(A)** Adult flies (5-7 days-old) subjected to ethyl butyrate (EB) along three concentrations (v/v). Data represent mean ± SEM with 350-400 individuals per groups and concentrations. When two different Aβ42 constructs were expressed in the GH298-Gal4 domain (genotypes: *GH298-Gal4/UAS-Aβ42* and *GH298-Gal4/UAS-Aβ42/UAS-GFP*^*nls*^), olfactory responses to 10^−3^ and 10^−2^ concentrations were consistently repulsive (full and dotted grey lines respectively) compared to *GH298-Gal4* controls (black line) and *GH298-Gal4/UAS-PI3K* flies (blue line). However, When PI3K and Aβ42 were co-expressed in GH298 neurons (genotype: *GH298-Gal4/UAS-PI3K/UAS-Aβ42*), the olfactory index returned to the normal profile (red line). Black and grey asterisks: comparison between *GH298-Gal4/UAS-Aβ42* and *GH298-Gal4/UAS-Aβ42/UAS-GFP*^*nls*^ flies with respect to control. ζ= \*\**p*<0.001, comparison between *GH298-Gal4/UAS-Aβ42* and *GH298-Gal4/UAS-PI3K/UAS-Aβ42* flies. This return to normal olfactory perception when PI3K and Aβ42 are simultaneously addressed to *GH298* neurons is also reproduced with IAA odorant (**Suppl. Fig. S1**). **(B)** Equivalent assay in the *krasavietz-Gal4* domain. The MB-Gal80 represses the Gal4 driver in the mushroom body neurons. Olfactory responses from *krasavietz-Gal4/MBGal80/UAS-Aβ42* flies were altered (full grey line) compared to *krasavietz-Gal4/MBGal80* control individuals (full black line). The simultaneous co-expression of *PI3K* and *Aβ42* restores the normal response to [10^−3^] and [10^−1^]. Note the olfactory response of theses flies at [10^−2^] above control values (red asterisk=\**p*<0.05). Black asterisks: comparisons between *krasavietz-Gal4/MBGal80/UAS-Aβ42* versus *krasavietz-Gal4/MB-Gal80* control flies. ζ=\*\*\**p*<0.0001, comparison between *krasavietz-Gal4/MBGal80/UAS-Aβ42* and *krasavietz-Gal4/MB-Gal80/UAS-PI3K/UAS-Aβ42*. \**p*<0.05; \*\**p*<0.001; \*\*\**p*<0.0001 (Student’s *t* test).

Gal4 expression domains often include cell subsets beyond those aimed in behavioral experiments, inhibitory local interneurons in the case of *GH298*, or are transiently expressed during development in other tissues (Casas-Tintó et al., 2017). To confirm the PI3K protective effect in other subset of olfactory neurons, we carried out a similar experiment using the driver *krasavietz*-Gal4 which targets a group of 5-8 excitatory local interneurons mainly. Due to the fact that the *krasavietz* domain also extends to extrinsic mushroom body neurons, we employed a *MB-Gal80* construct to silence the *krasavietz-Gal4* in these cells. The effects on EB perception were akin to those observed with the inhibitory local interneurons (**Fig. 4B**). Thus, we can conclude that the protective effects of PI3K upon the Aβ42-dependent toxicity are reproduced in olfactory neurons.

Aβ42 is also known to affect severely the organism lifespan when expressed throughout the nervous system, using a pan-neural *elav-Gal4* driver (Iijima et al., 2004). Therefore, we aimed to study here the overexpression of PI3K with the same adult-onset design. As expected, Aβ42 flies decreased their lifespan while PI3K increased it (**Suppl. Fig. S2A**). The joint co-expression, PI3K/Aβ42, significantly ameliorated the negative effect of Aβ42, albeit partially (**Suppl. Fig. S2A**).

Taken together, these data indicate that PI3K overexpression rescues Aβ42-dependent locomotor deficits and olfactory perception changes and ameliorates organism longevity defects. Thus, the PI3K-protected synapses seem functional throughout all neurons targeted here to the extent of improving the fly lifespan.

### 3.4 PI3K extends cell survival of human neurons exposed to Aβ42 oligomers

To validate if the results of PI3K in fly neurons affected by Aβ42 can be reproduced in human cells, we examined the effects of PI3K activation in human neuroblastoma SH-SY5Y cells treated with Aβ42 oligomers. Cells were cultured and differentiated with retinoic acid and 1% serum for 48 h prior to Aβ42 oligomers treatments. Aβ42 oligomers where added to the cell culture 72 h after cell seeding (48 h after differentiation), at 3 different concentrations (Kayed and Glabe, 2006). To stimulate PI3K activity we used the peptide PTD4-PI3KAc, previously proved to trigger PI3K activation both *in vitro* and *in vivo* (Cuesto et al., 2015, 2011; Enriquez-Barreto et al., 2014). PTD4-PI3KAc was administered at 3 different time points, before Aβ42 (pre-treatment), simultaneously with Aβ42 (co-treatment) or after Aβ42 addition (post-treatment) (**Fig. 5**). As expected, Aβ42 oligomers did not affect cell viability at low concentrations (18 μg/ml). However, the percentage of cells alive after the treatment was significantly decreased at 36 μg/ml or 72 μg/ml concentrations. These results confirmed the effectiveness of the oligomer preparation protocol and were used as positive internal controls in the experiment (**Fig. 5**).

**Figure 5.**
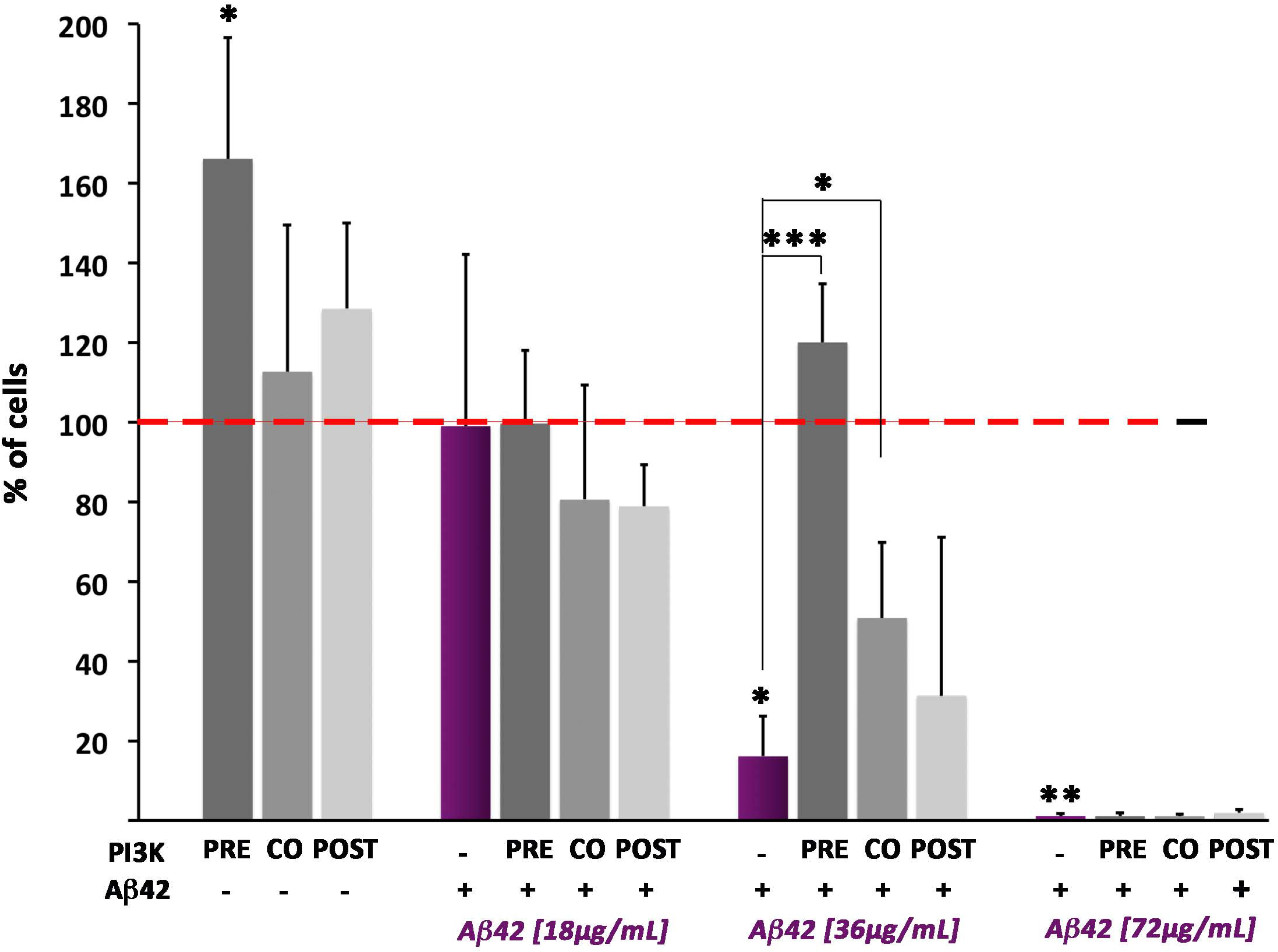
PI3K neuroprotection from Aβ42 effects is reproduced in human cells. Differentiated neuroblastoma SH-SY5Y human cells were treated with increasing concentrations of Aβ42 oligomers (18, 36 and 72 μg/ml) and PTD-PI3KAc peptide (50 μg/ml) administered 24 h prior (pre), at the same time (co) or 24 h after (post) the Aβ42 treatment. Data ± SD from Cell Titer Glo luminescence is shown as percentage of viability compared to control values of cells exposed neither to Aβ42 nor to PTD-PI3KAc (red-dotted line). At 32 μg/ml of Aβ42 treatment, the deleterious effects of the oligomers are evident and the PI3K protection is most effective if cultures are pre- or co-treated. At 72 μg/ml of Aβ42 treatment, the effects are so deleterious that no protection seems possible.

PTD4-PI3KAc peptide showed no cell viability effects in non-Aβ42-treated cells at 48 h or 72 h, but it significantly increased cell viability when administer 24 h after differentiation. This effect can be attributed to the brief period of starvation and differentiation that the cells had at this time (24 h) that could allow cells to respond to mitogen stimuli, like PI3K pathway effectors. More interestingly, PI3K increases cell viability only when PTD4-PI3KAc peptide is added 24 h before or simultaneously with Aβ42, at an oligomer concentration of 36 μg/ml **(Fig. 5)**. No protection was found at higher Aβ42 concentrations (72 μg/ml).

These data prove that PTD4-PI3KAc protects human neurons from Aβ42 oligomers toxic effects, validating the *in vivo* results obtained in flies. Presumably, the protective effect of this peptide will be elicited through the activation of the cell endogenous PI3K since the peptide has no kinase activity per se (see Mat & Meth.).

### 3.5 Protection from synaptic effects takes place through non-canonical PI3K signaling

Being PI3K a shared element in several signaling pathways (Acebes and Morales, 2012), we explored two possibilities, 1) the insulin receptor canonical PI3K/Akt pathway, which is known to elicit pro-survival and cell growth responses (Basu et al., 2012; Dyer et al., 2016), and 2) the recently revealed pro-synaptogenesis pathway (Jordán-Álvarez et al., 2017). To assay the effect of the classical pathway effector, we overexpressed *Target of Rapamycin* (*mTOR*), a downstream factor of PI3K, and we analyzed the synaptic signal in abdominal motor neurons of 15 days-old adults. The quantification of the synaptic BRP signal indicates that mTOR does not protect neurons from Aβ42 toxicity (**Fig. 6A,B-D,E,G**). This result is consistent with previous reports in *Drosophila* and mammals where mTOR is ineffective to change synapse number (Cuesto et al., 2011; Martín-Peña et al., 2006). On the other hand, Medea, a member of the pro-synaptogenesis pathway acting downstream from PI3K (Jordán-Álvarez et al., 2017), fully prevents the deleterious effects of Aβ42 on synapses when downregulated (**Fig. 6C,F,G**).

**Figure 6.**
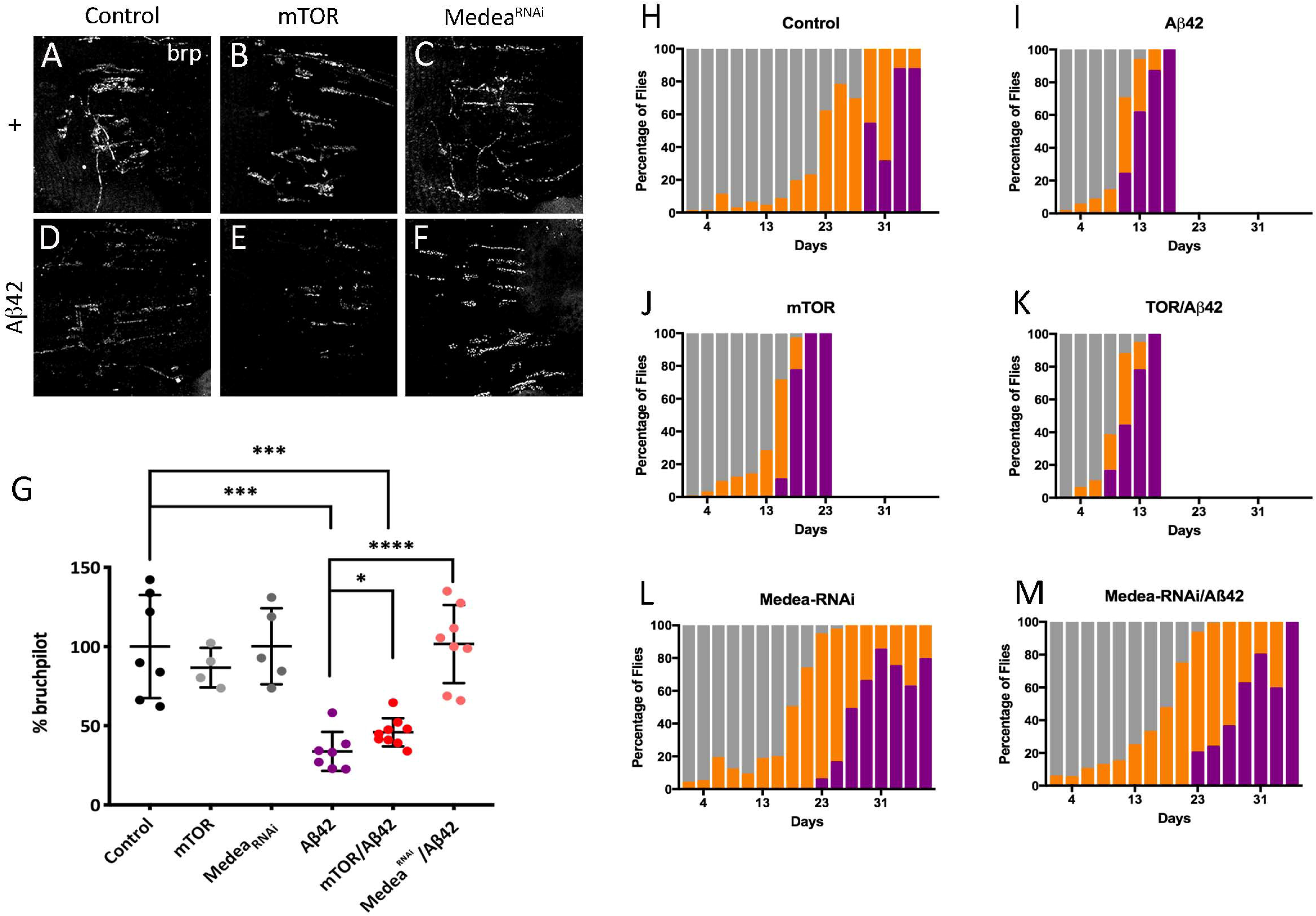
Suppression of Aβ42 effects is mediated through the synaptogenic PI3K pathway. **(A-F)** Representative confocal images of 15 days adult females NMJ corresponding to the ventral muscle in the third abdominal segment. Active zones are revealed by nc82 antibody. **(G)** Quantification of synaptic area at 15 days. **(H-M)** Climb assay with color-coded performance following the same protocol and criteria as in Fig. 2. Note the severe enhancement of Aβ42 defects by mTOR, not a member of the synaptogenesis PI3K signaling, while Medea^RNAi^, a member of that pathway, does suppress them fully. Data represent mean ± SD. Note the neuroprotection elicited by Medea^RNAi^ but not by mTOR. Numerical data are shown in **Supp. Table S1**.

To test the behavioral outcome of the synapse loss prevention by Medea down-expression, we performed climbing experiments using the same genotypes tested for synapse quantification. mTOR/Aβ42 flies showed the same reduction in climbing activity as either of both factors separately (**Fig. 6H-K**). Non-climbing flies begun to be detected at day 9 (Aβ42) or day 15 (mTOR), while, in controls, this class became evident from day 29 onwards. By contrast, the down-regulation of Medea rescued the climbing phenotype of Aβ42 to a large extent (**Fig. 6G,L,M**).

Knowing that PI3K ameliorates lifespan in Aβ42 expressing flies, we wondered if mTOR overexpression or Medea down-expression could also affect longevity. Lifespan studies showed a significant decrease in mTOR/Aβ42 flies, as well as in mTOR overexpression alone (**Suppl. Fig. S2B**). Actually, mTOR overexpression aggravated the lifespan reduction caused by Aβ42 alone, while downregulating Medea lead to a normal lifespan in Aβ42 expressing flies. Thus, PI3K overexpression and Medea attenuation, two manipulations previously shown to upregulate synapse number, effectively decrease Aβ42 synaptotoxicity, allowing a better locomotor performance and increasing fly lifespan. Taken together, these data indicate that the protective effects of PI3K over Aβ42 toxicity rely on the synaptogenic pathway signaling, rather than on the canonical insulin receptor pathway.

### 3.6 PI3K increases Aβ42 insoluble aggregates in Drosophila and human neurons

To understand the causative effects of PI3K over Aβ42 accumulation, we analyzed total Aβ42 accumulation in 15 days-old fly brains by immunohistochemistry. Aβ42-overexpressing adult brains showed a noticeable 6E10 positive signal, which was absent in control or PI3K-overexpressing brains. Surprisingly, this positive signal was increased in intensity and extent in PI3K/Aβ42 brains (**Fig. 7A-D**). To confirm this result, same aged adult head homogenates were analyzed by Western blot (**Fig. 7E**). Total Aβ42 protein levels were 2.5-fold higher in PI3K/Aβ42 compared to Aβ42 alone. To rule out a putative effect on transcription, qRT-PCR was performed with same aged adult head homogenates and the results showed no changes in total Aβ42 mRNA, discarding this possibility (**Fig. 7F**). Thus, mechanistically, PI3K protects from Aβ42 toxicity and elicits an increase of the amyloid protein deposits but does not affect the transcription of the Aβ42 construct.

**Figure 7.**
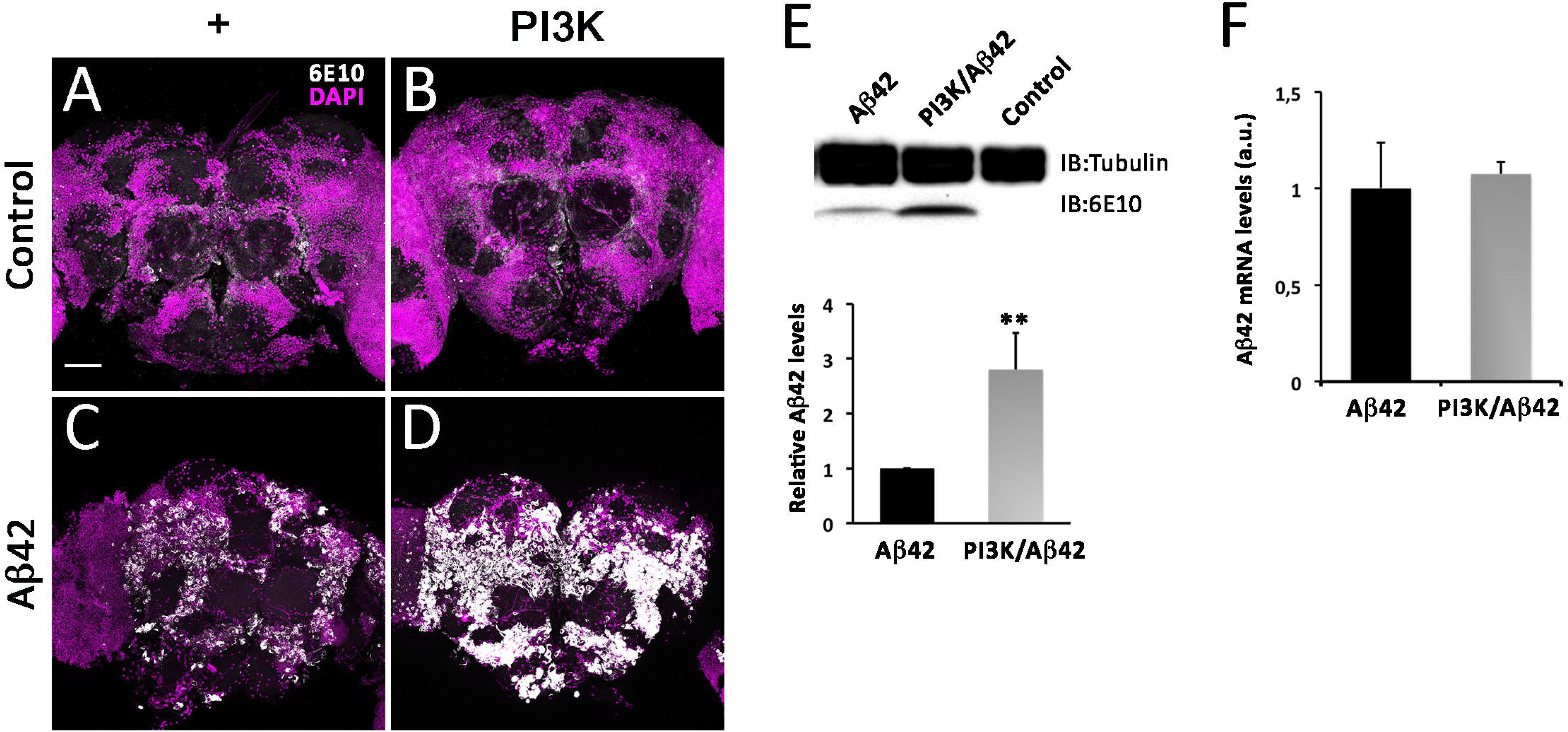
PI3K activation increases Aβ42 protein deposits without affecting transcription. **(A-D)** Representative confocal images of 15 days old adult heads stained with 6E10 antibody of the indicated genotypes. Note the large increase of amyloid deposits (white) revealed by the 6E10 antibody. Cell nuclei are marked by DAPI (magenta). **(E)** Western blot of adult heads of the same genotypes and age. Data correspond to three independent Western blots normalized for Tubulin content as internal control. Note the significant increase of the 6E10 signal elicited by PI3K activation. **(F)** Quantitative RT-PCR data from the same genotypes and ages. All data are ± SD. Scale bar is 50μm.

Since the various amyloid aggregates of Aβ42 can evolve different levels of toxicity, we tested the nature of the Aβ42 positive deposits by the criterion of Thioflavin-S staining. This dye binds specifically to proteins with β-sheet conformation and labels Aβ42 fibers and plaques, constituents of the insoluble fraction of β-amyloid aggregates. Quantification of the positive Thioflavin-S signal in 7 and 15 days-old brains demonstrated a significant increase in the insoluble fraction of Aβ42 deposits at 15 days of age in the PI3K/Aβ42 brains compared to age-matched Aβ42 alone (**Fig. 8A,B**). This result was confirmed by a 4-fold increase of the Aβ42 fraction in Western-blot of insoluble protein extracts from 15 days-old adult heads (**Fig. 8C**). These data show that PI3K triggers a change in Aβ42 aggregation into a more insoluble species.

**Figure 8.**
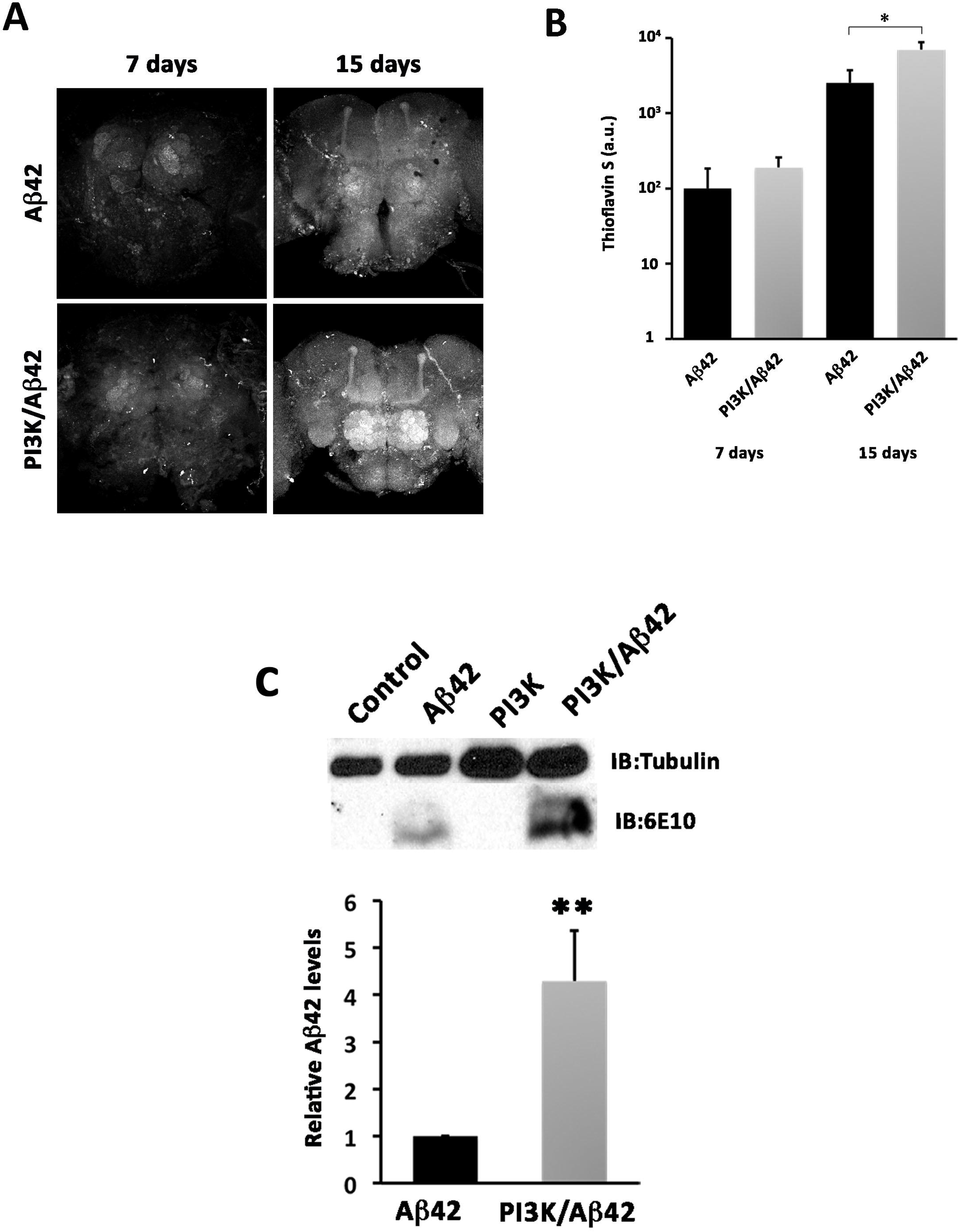
PI3K increases amyloid Aβ42 insoluble fraction. **(A)** Representative confocal images of 7 and 15 days old adult brains stained with Thioflavin. Scale bar is50μm. **(B)** Quantification of Thioflavin signal. Note the progressive signal increment with age and, in particular, with PI3K co-expression. **(C)** Immunoblot showing insoluble Aβ levels in control, PI3K, Aβ42 and Aβ42/PI3K adult heads of 15 days-old flies. Histograms represent mean Aβ levels normalized to Tubulin on three independent blots.

Beyond the effects in fly neurons, we tested whether human neurons could also exhibit the PI3K-induced increase of Aβ42 aggregation deposits. SH-SY5Y cell-line human neuroblastoma cells, were treated with Aβ42 oligomers (36 μg/ml) and PTD4-PI3KAc peptide (50 μg/ml) following differentiation. As expected, no 6E10 positive deposits were found in cells treated either with vehicles or with PTD4-PI3K alone (**Fig. 9**). For practical reasons only Aβ42 and PTD4-PI3KAc/Aβ42 experiments were compared. Treatment with PTD4-PI3KAc in combination with Aβ42 oligomers produced a significant decrease in the total number of deposits compared to those found in Aβ42 treatment (**Fig. 9**). The deposits, however, were up to 6-fold larger in cell cultures co-treated with Aβ42 oligomers and the PI3K-activating peptide.

**Figure 9.**
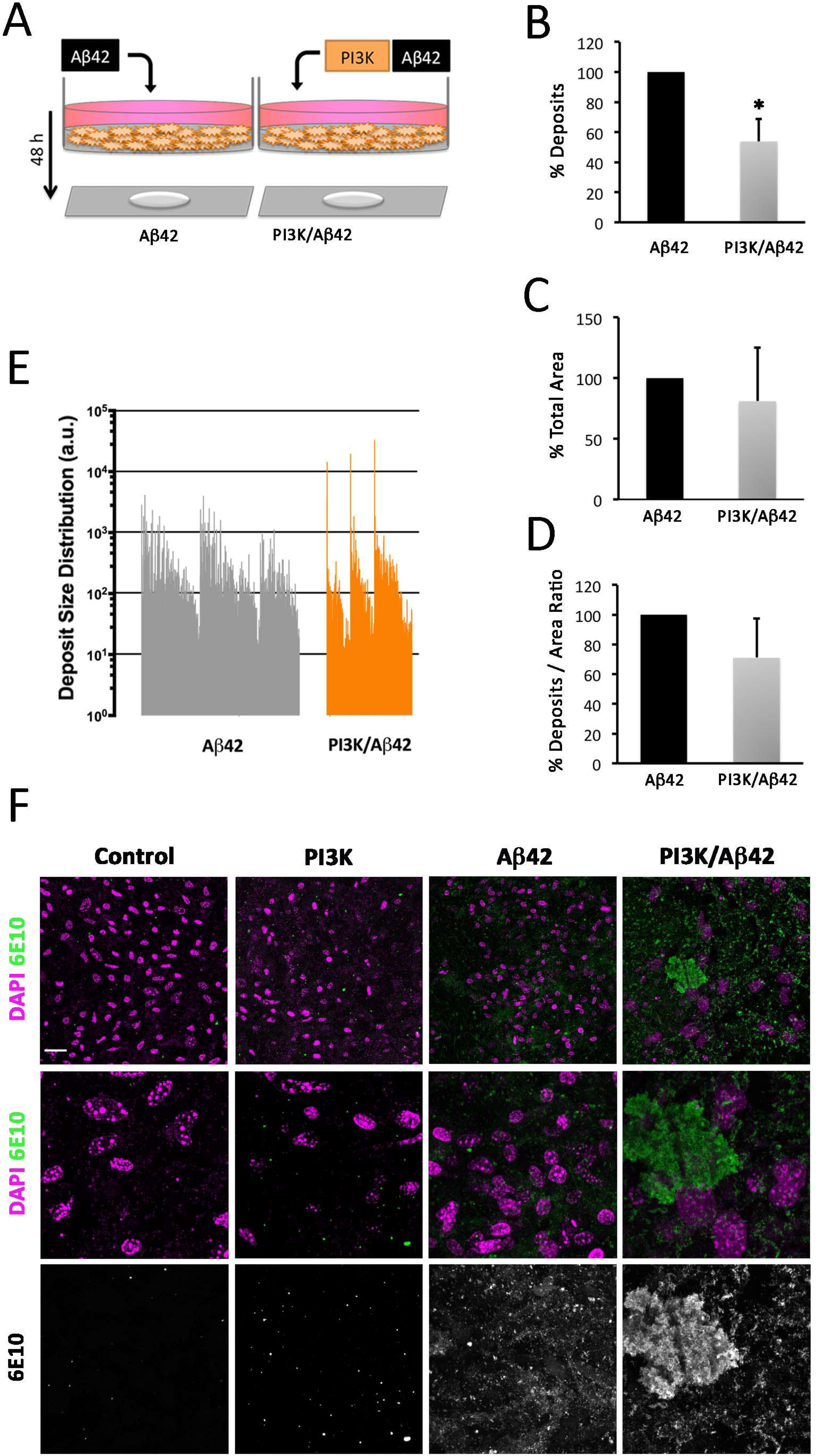
PI3K increases amyloid Aβ42 insoluble fraction in human cells. **(A)** Cartoon representing the experimental design in SHT-SY5Y cells differentiated and treated with Aβ42 oligomers (36 μg/ml) and PTD4-PI3KAc peptide (50 μg/ml) for 48 h. **(B-E)** Histograms representing total deposits **(B)**, size distribution **(C)**, total area **(D)**, and deposits to area ratio **(E)** in the cultures. **(F)** Representative confocal images of cell cultures showing Aβ (6E10) (green) and DAPI (magenta) signals. Note the large amyloid deposits in PI3K/Aβ42. Scale bar is 10 μm.

To further analyze the aggregation effect of Aβ42 oligomers, we questioned if these aggregates could occur in the extracellular space, where the majority of deposits are usually found. Four different experiments were designed: 1) an internal control to test Aβ42 oligomers aggregation, 2) cells treated with Aβ42 oligomers previously treated with PTD4-PI3KAc (pre-treatment), 3) cells treated with Aβ42 oligomers and PTD4-PI3KAc at the same time (co-treatment), and 4) cells treated with Aβ42 oligomers later treated with PTD4-PI3KAc (post-treatment). Supernatants were recovered and deposits were stained with 6E10 antibody (**Suppl. Fig. S3**).

The internal control of Aβ42 oligomers showed no 6E10 positive deposits, in the given time of 48 h. We found no significant differences when comparing neither the total number of deposits in the PTD4-PI3K pre-treatment, co-treatment and post-treatment experiments nor in the total deposit area (**Suppl. Fig. S3C,D**). However, the three treatments showed significant differences when the individual size of each deposit was analyzed (**Suppl. Fig. S3B**). PTD4-PI3K co-treatment had up to a 10-fold increase in deposit size, when compared to PTD4-PI3KAc pre-treatment and post-treatment. PTD4-PI3KAc post-treatment showed the smallest deposits among the three experiments of the study (**Suppl. Fig. S3E**).

These data indicate that the effects in Aβ42 aggregation elicited by PI3K in adult flies are reproduced in human SH-SY5Y differentiated neurons, which can occur in the extracellular space.

### 3.7 PI3K increases phosphorylation of Aβ42

Building on previous findings pointing to phosphorylation as a factor affecting Aβ42 seeding in the initial phase of the plaque formation (Kumar et al., 2016, 2011; Kumar and Walter, 2011), we addressed whether the PI3K kinase activity could play a role in the mechanism of Aβ42 aggregation. To explore this hypothesis, we applied the NetPhos 2.0 Server prediction software (www.cbs.dtu.dk) and identified three putative residues: pSer8 (Score: 0.963), pTyr10 (Score: 0.870) and pSer26 (Score: 0.787) as candidates (**Fig. 10A**).

**Figure 10.**
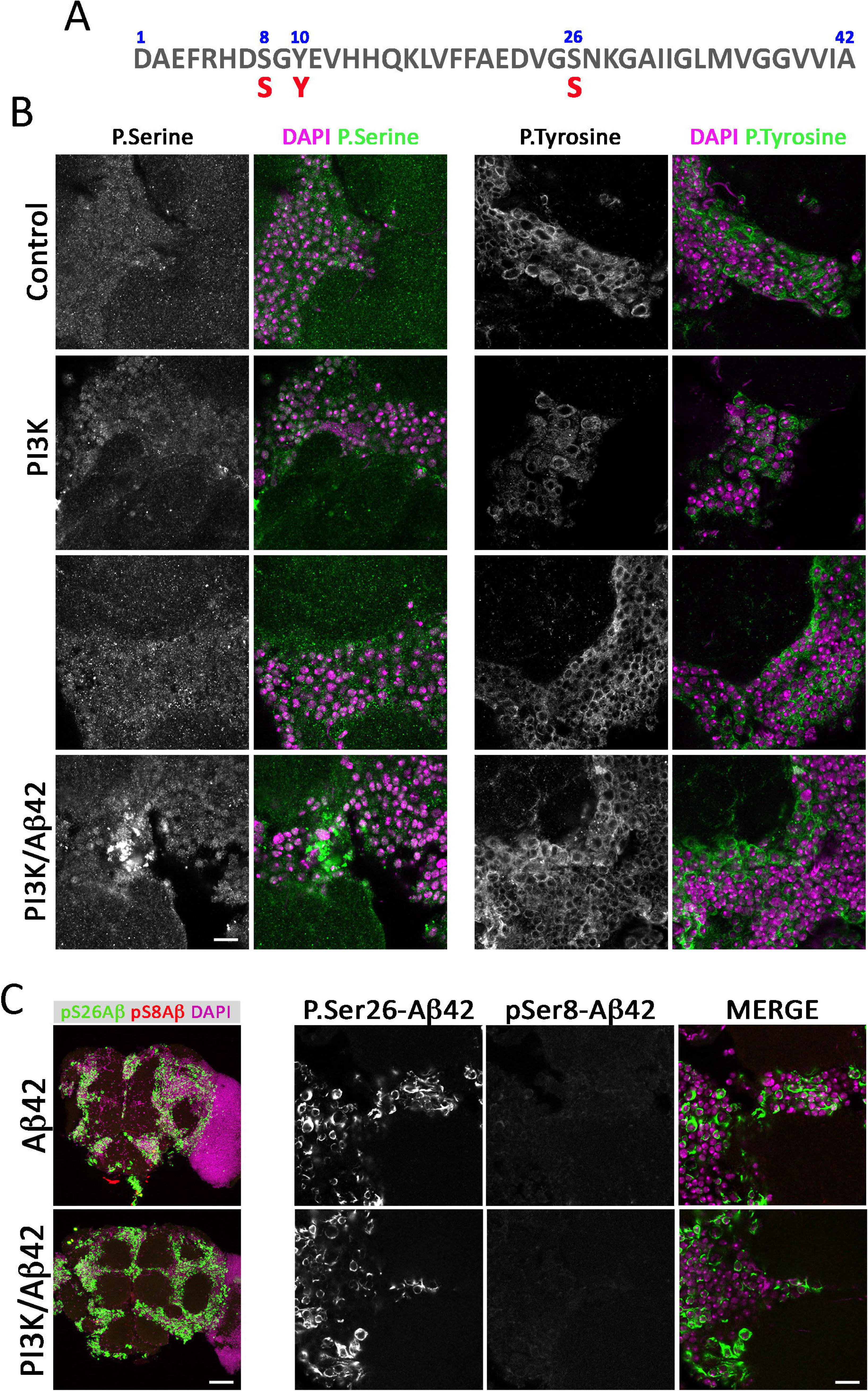
Aβ42 residues potentially targeted by PI3K. **(A)** Serine and Tyrosine residues in Aβ42 sequence subject to phosphorylation. **(B)** Images of p-Ser (green) and DAPI (magenta) stainings in 15 days old adult brains of the corresponding genotypes (see Figure 1 legend). Equivalent images of brains stained for p-Tyr (green) are shown in the right panels. Note the increment of p-Ser, but not p-Tyr, immunosignal in PI3K/Aβ42 brains. Scale bar is 10 μm. **(C)** Images of p-Ser26 (green), p-Ser8 (red) and DAPI (magenta) immuno-stainings in 15 days old adult brains of the corresponding genotypes. Scale bar is 10 μm **(B)** and 50 μm **(C)**.

Specific antibodies against phospho-Tyr and phospho-Ser were applied in 15 days-old brains in the four genotypes under study. The phospho-Ser signal was only incremented in PI3K/Aβ42 brains and had a deposit-like morphology (**Fig. 10B**). By contrast, phospho-Tyr signal was comparable in all genotypes and no deposit-like morphology was found (**Fig. 10B**). Co-localization of 6E10 and both phospho-Ser and phospho-Tyr was not feasible due to the requirement of formic acid for 6E10 immunostaining, which strongly alters phospho-peptides and membrane stability.

Aiming to discriminate between the two Serine residues, Ser-8 and Ser-26, as candidates for phosphorylation, we tested pSer-8-Aβ42 (1E4E11) and pSer-26-Aβ42 (7H3D6) antibodies (Kumar et al., 2016, 2013). No changes were found in pSer8-Aβ42 immunostainings in Aβ42 or PI3K/Aβ42 genotypes (**Fig. 10C**). Phospho-Ser26-Aβ42 immunostainings showed deposits with positive signal in both Aβ42 and PI3K/Aβ42. Unfortunately, the pSer-26-Aβ42 antibody seems to detect both phosphorylated and non-phosphorylated forms of Aβ42.

The data prompted us to conclude that PI3K overexpression in an Aβ42 background drives the accumulation of phospho-Serine positive deposits.

## 4. Discussion

Synapse loss is one of the first steps in the neurodegenerative process in Alzheimer’s disease. Previous *in vitro* and *in vivo* studies have demonstrated the age independent synaptogenic actions of PI3K (Acebes et al., 2011; Cuesto et al., 2011; Martín-Peña et al., 2006). Thus, we set out to study the potential effects of PI3K in ameliorating the AD features in an adult-onset *Drosophila* model. Benefitting from the temporal control of the Gal4/UAS binary system, we have driven the expression of human Aβ42 peptide in the adult central nervous system and followed the progression of the synaptic toxicity. Although another study has used a similar strategy based in the GeneSwitch system by feeding mifepristone (RU486) (Sofola et al., 2010), most AD models in *Drosophila* and other organisms, follow constitutive expression of the Aβ peptide, APP, PS1, tau, in their mutated or wild type forms (Ashe and Zahs, 2010; De Felice and Munoz, 2016; Iijima-Ando and Iijima, 2010; Wisniewski and Sigurdsson, 2010).

### 4.1 PI3K mechanisms in Aβ42-induced synapse toxicity

Protein kinases (PK) in general, can operate inside or outside the cell, where they can phosphorylate cell-surface proteins or extracellular substrates (Redegeld et al., 1999; Shaltiel et al., 1993; Walter et al., 1994). These Ecto- and Exo-PKs can use extracellular ATP, which is present in the brain at nanomolar concentrations, and can also increase locally upon certain stimuli (el-Moatassim et al., 1992; Gourine et al., 2007; Melani et al., 2005). The extracellular Aβ42 deposits identified in this study could, thus, be originated directly outside of the cell or within the presynaptic side, where they are genetically co-expressed, followed by their extrusion.

In turn, PTEN is a lipid phosphatase that inhibits PI3K activation (Maehama and Dixon, 1998), and it mediates Aβ42 synaptic and cognitive impairments (Knafo et al., 2016). PTEN can regulate LTD (Jurado et al., 2010) and it can be targeted by Aβ42 in synaptic spines. Furthermore, pharmacological administration of a PTEN membrane-permeable peptide, unable to interact with Aβ42, rescues synapse deterioration (Knafo et al., 2016). Other PI3K regulators, like the signaling inhibitor GSK3-β, are known to elicit anti-synaptogenic actions in *Drosophila* and mice (Cuesto et al., 2015; Martín-Peña et al., 2006), with concomitant effects in LTP and long-term memory (Hooper et al., 2008). In addition, Jun kinase/AP-1 and Wnt signaling are also members of the synaptogenic pathway, as they are regulated by GSK3-β (Franciscovich et al., 2008). In AD, GSK3-β is known to phosphorylate Tau and regulate microtubule dynamics (reviewed in (Llorens-Martín et al., 2014). PIP3, synthetized by PI3K, regulates PSD-95 clustering and synaptic AMPA receptors in spines, affecting synaptic strength (Arendt et al., 2010). All these features compose an scenario in which PI3K is situated in an strategic position to influence opposite pro- and anti-synaptogenic signaling, whose equilibrium ultimately determines the number of synapses that a neuron stablishes (Jordán-Álvarez et al., 2017).

Transcriptional changes do not seem to be part of the mechanism elicited by PI3K synaptogenic effect. Our data suggest that neither Aβ42 nor PI3K promote mRNA changes in three major AZ components (*bruchpilot*, *liprin* and *synaptobrevin* genes). Nonetheless, other transcriptional alterations, or even post-transcriptional regulations triggered by PI3K could modify synaptic proteins availability, turnover, degradation and/or trafficking, leading to the synaptic changes described here.

Microtubule dynamics showed that Aβ42 reduces EB1 microtubule binding protein comets density, and that this effect was also suppressed by PI3K activation. Defects in microtubule dynamics alter protein trafficking along the axon (Guedes-Dias et al., 2019). Brp, as well as other components of the AZs and PSDs, need to be transported via microtubules, and decreased microtubule stability can hamper, not only the formation of new AZs or PSDs structures, but also synaptic protein turnover. Aβ42 effects on microtubule dynamics might explain the reduction in the synaptic area in adult NMJs that did not depend on transcriptional changes. How PI3K suppresses the Aβ42-dependent impairment of microtubule dynamics has not been addressed in this study. However, among the wide diversity of PI3K-dependent signaling pathways, some are linked to actin cytoskeleton via Rac activation and are Akt-independent (Lien et al., 2017), while others connect microtubules with insulin receptors via GSK3-β (Chiu et al., 2008). If equivalent pathways would exist in *Drosophila*, they could support the observed suppression of defective microtubule dynamics by PI3K.

Aiming to discriminate among PI3K effector pathways, we assayed here mTOR and Medea as potential mediators of Aβ42. Only the latter proved to be effective indicating that, in the Aβ42 context, PI3K proceeds through the Medea signaling. In addition, Medea downregulation is known to attenuate BMP signaling (Wisotzkey et al., 1998) which is altered in AD (Crews et al., 2010; Kang et al., 2014). On the contrary, mTOR appears to enhance the neurotoxic effects of Aβ42 in lifespan and reduces synapse number, similarly to Aβ42 flies when expressed alone (**Supp. Fig. S2**). This result is in line with others, where mTOR is found to participate in memory formation and consolidation, and seems targeted by Aβ42 (Lin et al., 2014; Mueed et al., 2019; Uddin et al., 2018). Thus, mTOR inhibition may be another experimental strategy, independent from PI3K activation, to be considered in future AD studies.

### 4.2 PI3K mechanisms in Aβ42 dynamics

Aberrant protein phosphorylation has been described in the brain of AD patients (Chung, 2009). Here, we identified a new role of PI3K in Aβ42 accumulation with *in vivo* and *in vitro* results that show that PI3K activation induces aggregation of Aβ42, leading to insoluble, less toxic, aggregates. Previous studies consider insoluble deposits of Aβ42 as rather benign species (DaRocha-Souto et al., 2011; Moreth et al., 2013b), in contrast to soluble Aβ42 oligomers. Although insoluble deposits would seem refractory to degradation, at least three enzymes are known to degrade Aβ42, Neprilysin, Insulin-degrading enzyme (IDE) and Endothelium-converting enzyme (ECE) (Grimm et al., 2013; Turner et al., 2004).

Post-translationally modified Aβ species have been identified (Kummer and Heneka, 2014; Kuo et al., 2001) including truncation (Härtig et al., 2010), racemization (Murakami et al., 2008), isomerization (Shimizu et al., 2000), pyroglutamination (Kuo et al., 1997), metal-induced oxidation (Dong and Bai, 2003) and phosphorylation (Kumar et al., 2011; Milton, 2001). Particularly, phosphorylation can be potentially achieved at three different Aβ42 residues, Ser-8, Ser-26, and Tyr-10. In addition, Aβ is reported to undergo phosphorylation by protein Kinase A (PKA) and by Cyclin-dependent kinase 2 (CDC-2) *in vitro* (Milton, 2001). Phosphorylation at Ser-8 by PKA was observed in free extracellular Aβ rather than in its precursors (APP or β-CTF) and promoted the formation of toxic aggregates. Also, phosphorylated deposits from APP transgenic mice were found concentrated in the center of the plaque (Kumar et al., 2011). Phosphorylation in Ser-8 has been documented to induce Aβ resistance to degradation by Insulin-degrading enzyme (Kumar et al., 2012).

One study described Aβ phosphorylation at Ser-26 by CDC-2 *in vitro* (Milton, 2001) whereas another showed that pSer26-Aβ42 is abundant in intra-neuronal deposits at very early stages of AD (Kumar et al., 2016). However, potential changes in Aβ42 aggregation were not investigated. Nevertheless, cell treatment with Olomoucine prevents cytotoxicity and Aβ phosphorylation, suggesting that aggregative forms favored by Ser-26 phosphorylation enhance Aβ42 progressive toxicity. In contrast, our data suggest that PI3K-induced phosphorylation in Ser-26 ameliorates Aβ42 effects and promotes insoluble Aβ42 assembly formation. A recent *in vitro* analysis of Aβ40 phosphorylated at Ser-26 describes that this modification impairs fibrillization while stabilizing monomers and non-toxic soluble aggregates. Computational studies predicted that a phosphate-group at Ser-26 could rigidify the region and interfere with a fibril-specific salt-bridge (Rezaei-Ghaleh et al., 2014).

### 4.3 Structural considerations and perspectives

A specific region in the Aβ sequence comprising residues 25-29 generates a bend-like structure that juxtaposes the hydrophobic faces of the two cross-β units. This bend is important for Aβ pathogenic aggregation (Fawzi et al., 2008; Grant et al., 2007; Murray et al., 2009). Other studies have further emphasized the importance of the Gly-25–Ser-26 dipeptide in Aβ42 monomer structure by finding changes in aggregative properties upon biochemical modifications in these two residues (Roychaudhuri et al., 2014). The potential use of drugs that could alter the interactions of this dipeptide with neighboring side-chain atoms, or with the core sequence of the peptide, has been proposed.

Moreover, specific mutations in APP, such as Arctic (E22G), Italian (E22K), Dutch (E22Q), Osaka (E22Δ) and Iowa (D23N), are known to induce aggregation. These variants can cause early onset familial AD (Benilova et al., 2012). Thus, given the biochemical and biophysical properties of Aβ peptide, and the evidences reported here, new strategies based on specific residue modifications like phosphorylation or dephosphorylation could be explored.

The suppression mechanism of neurotoxicity elicited by PI3K activation is very different from others, such as Lithium treatment, where the inhibition of translation seems to be the targeted step (Sofola-Adesakin et al., 2014). The seemingly counter-intuitive observation that increasing Aβ deposits is neuroprotective must be evaluated in the context of the equilibrium between soluble, albeit more toxic, oligomers and the relatively innocuous insoluble aggregates. PI3K seems to displace the equilibrium towards the insoluble aggregate forms. Given the conserved nature of PI3K pathways (Ruggero and Sonenberg, 2005), including the synaptogenesis signaling (Cuesto et al., 2011; Enriquez-Barreto et al., 2014; Jordán-Álvarez et al., 2017), the data reported here invite to consider PI3K activation as a suitable candidate to prevent AD progression. The PTD4-PI3KAc or equivalent peptides seem a powerful tool in this context.

## Acknowledgements

We appreciate fly strains from Bloomington Stock Center and the VDRC repository. We would like to thank Prof. David Van Vactor (Harvard University) for hosting M.A. during a working stay to perform the live imaging experiments on microtubule dynamics. We also appreciate the help of Dr. Cristina Martin-Higueras for paper editing. Research was funded by grant BFU2015-65685-P from the Spanish Ministry of Economy.

## Supplementary Data

**Supplementary Figure S1.- PI3K prevents Aβ42-induced olfactory perception defects to isoamyl acetate.** Adult 5-7 days aged flies were subjected to an olfactory assay for isoamyl acetate (IAA) with three concentrations (v/v). The expression of two different Aβ42 constructs in the *GH298-Gal4* domain (genotypes: *GH298-Gal4/UAS-Aβ42* and *GH298-Gal4/UAS-Aβ42*/*UAS-GFP*^*nls*^, full and dotted grey lines respectively) yields more repulsive responses to [10^−3^] and [10^−2^] compared to *GH298-Gal4* controls (full black line) and *GH298-Gal4/UAS-PI3K* flies (dotted black line). However, when PI3K and Aβ42 were co-expressed in GH298 neurons (genotype: *GH298-Gal4/UAS-PI3K/UAS-Aβ42*), the olfactory index returned to the normal profile (red line). Black asterisks: comparisons between *GH298-Gal4/UAS-Aβ42* and *GH298-* with respect to control. Red asterisks: comparisons between *GH298-Gal4/UAS-Aβ42* and *GH298-Gal4/UAS-PI3K/UAS-Aβ42* flies. **p<0.001; ***p<0.0001 (Unpaired Student’s t test with Welch’s correction).

**Supplementary Figure S2.- Aβ42-induced reduction of lifespan is prevented by PI3K synaptogenesis signaling. (A)** Longevity profiles of color coded genotypes at 29°C. PI3K activation suppresses the lifespan reduction caused by Aβ42 to a large extent. Numerical data are shown in **Supp. Table S1**. **(B)** Equivalent profiles for other genotypes including factors that do not participate in the PI3K synaptogenic pathway, mTOR, or included in that signaling, Medea^RNAi^. Note the full suppression by the later factor and the enhanced lifespan reduction caused by mTOR. Numerical data are shown in **Supp. Table S1**.

**Supplementary Figure S3.- PI3K-induced increase in Aβ42 aggregation can be reproduced outside the cell. (A)** Cartoon representing the experimental design with SHT-SY5Y differentiated cells treated with Aβ42 oligomers (36μg/ml) and PTD-PI3KAc peptide (50μg/ml) at different times of administration. Supernatants were stained with 6E10 antibody. Cells treated only with Aβ42 oligomers were used as control. **(B-E)** Histograms representing percentage of deposits/area ratio **(B)**, percentage of deposits **(C)**, percentage total area **(D)**, and deposit size distribution **(E)** of Aβ42 and PI3K/Aβ42 treated cells. Control cells treated only with Aβ42 oligomers did not develop recognizable 6E10 positive deposits.

**Supplementary Table S1.-** Data from the climbing and life span assays with the corresponding statistical tests. Climbing assay performance is color coded as in main Figures 2 and 6.

