## Supplementary figures and images for "PI3K activation prevents Aβ42-induced synapse loss and favors insoluble amyloid deposits formation"

### Supp Fig S1

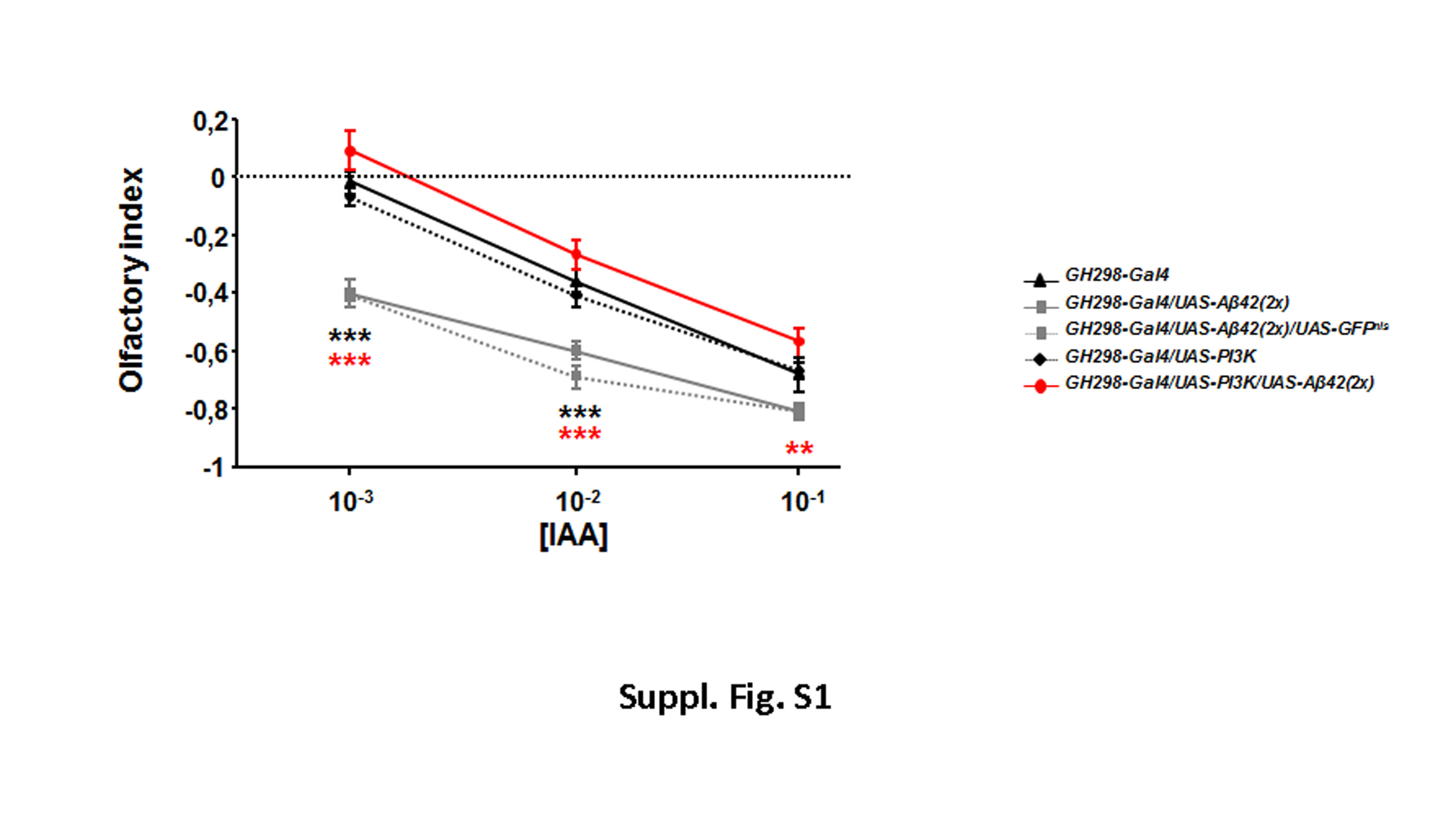

### Supp Fig S2

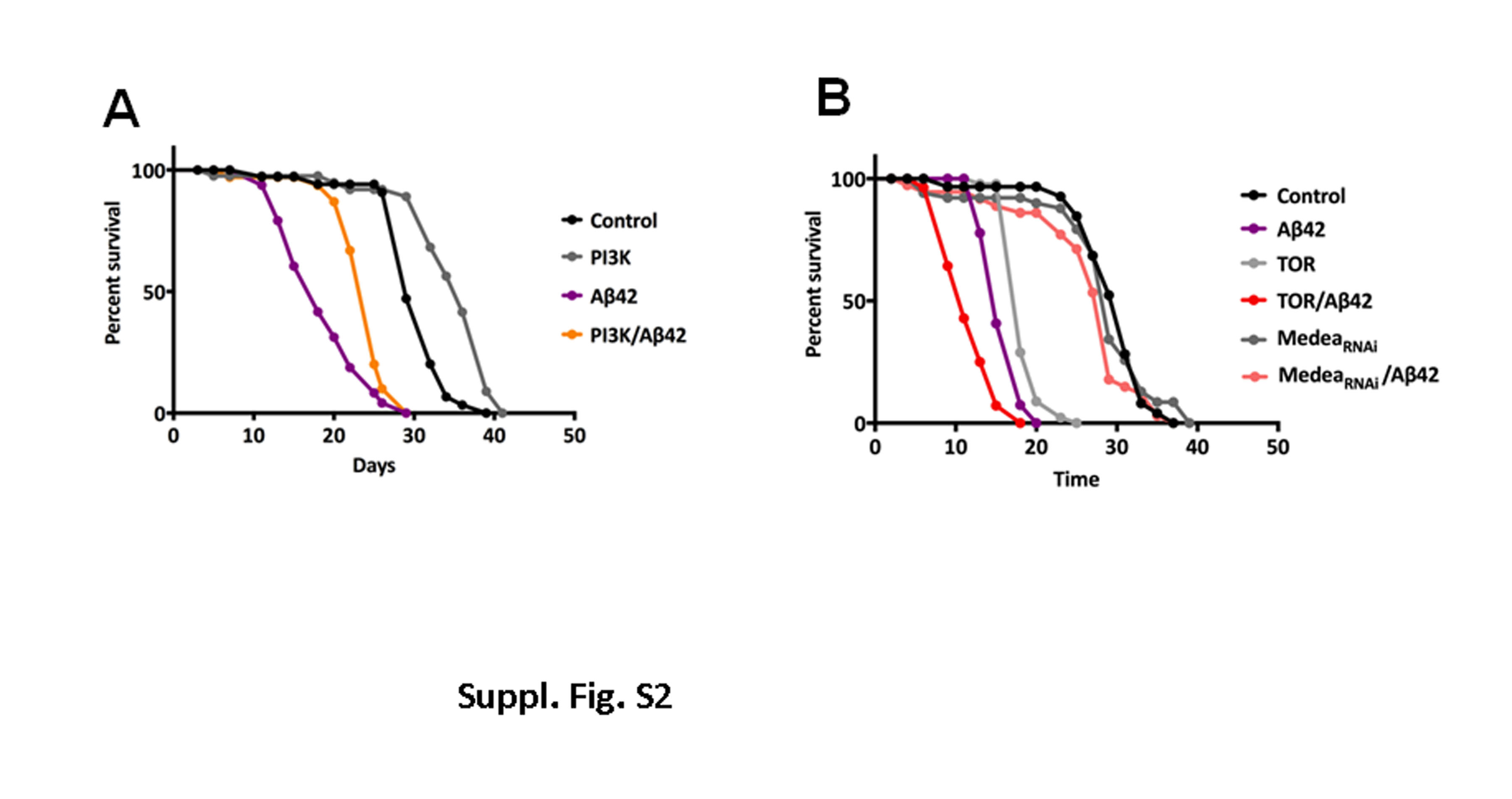

### Suppl Fig S3

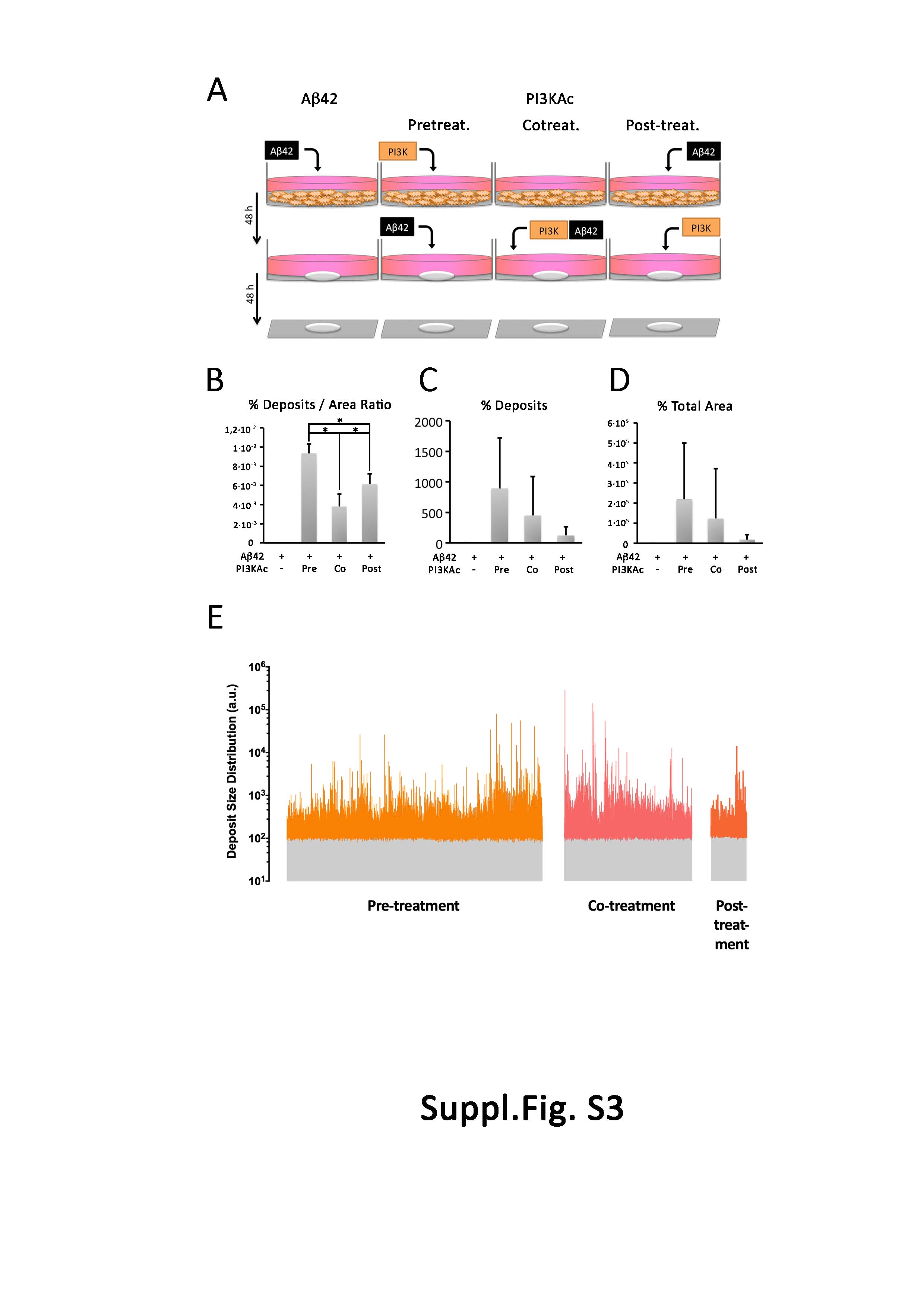

### Suppl Table1

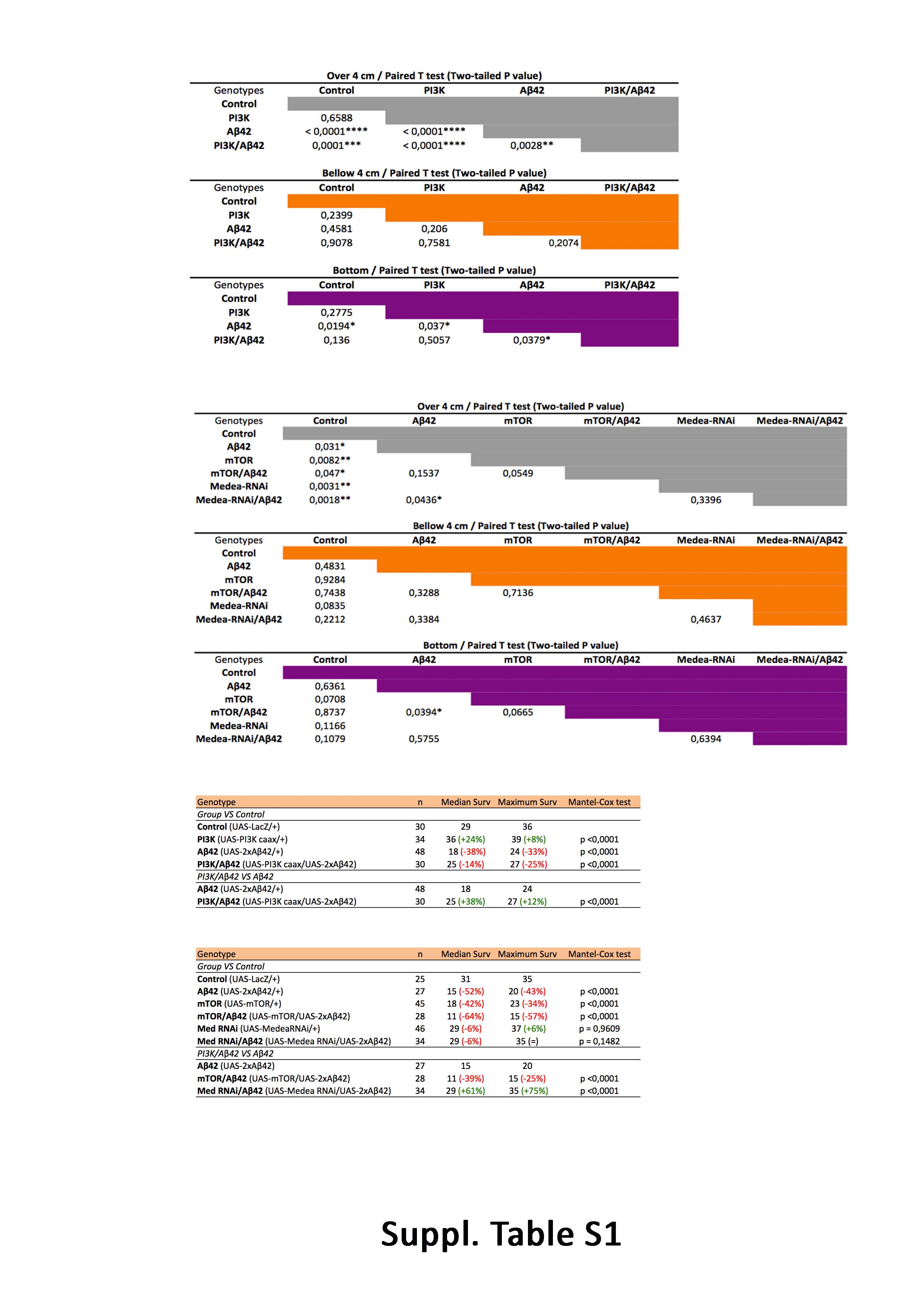
